# A computational model to unravel the function of amyloid-*β* peptide in contact with a phospholipid membrane

**DOI:** 10.1101/2020.01.17.910182

**Authors:** Pham Dinh Quoc Huy, Pawel Krupa, Hoang Linh Nguyen, Giovanni La Penna, Mai Suan Li

## Abstract

The normal synapse activity involves the release of copper and other divalent cations in the synaptic region. These ions have a strong impact on the membrane properties, especially when the membrane has charged groups, like it is the case of synapse. In this work we use an atomistic computational model of dimyristoyl-phosphatidylcholine (DMPC) membrane bilayer. We perturb this model with a simple model of divalent cation (Mg^2+^), and with a single amyloid-*β* (A*β*) peptide of 42 residues, both with and without a single Cu^2+^ ion bound to the N-terminus. In agreement with experimental results reported in the literature, the model confirms that divalent cations locally destabilize the DMPC membrane bilayer, and, for the first time, that the monomeric form of A*β* helps in avoiding the interactions between divalent cations and DMPC, preventing significant effects on the DMPC bilayer properties. These results are discussed in the frame of a protective role of diluted A*β* peptide floating in the synaptic region.

**Author summary:** We modelled the behavior of a Mg^2+^ divalent cation, with the size of Zn^2+^ and Cu^2+^, in contact with a phosphatidyl lipid bilayer. We also modelled the monomeric amyloid-*β* peptide 1-42, both free and Cu-loaded, the latter mimicking the final step of the binding between the peptide and the divalent cation. On the basis of the simulation results, we propose that the peptide hinders the strong interactions between the divalent cation and the membrane.

## Introduction

Alzheimer’s disease is a degenerative disease, with one histological hallmark being extracellular deposits in the central nervous system [1]. These deposits are made of amyloid peptides originated by the amyloid precursor protein (APP), a trans-membrane protein with a multimodal function [2]. Amyloid-*β* (A*β*) peptides are produced with proteolysis of APP at the membrane interface, by the enzymes *β* and *γ*-secretases. The *γ* cleavage, that produces most of the neurotoxic peptides (39-42 residues), occurs deeper in the membrane bilayer compared to *β* cleavage [3]. The production of these toxic peptides at the membrane interface can have many important implications [4], even before peptide aggregation occur and when oligomers are more abundant than protofibrils [5, 6]: i) the toxic pathway can be influenced by interactions between peptides and of peptides with the membrane; ii) the peptide, depending on its concentration, can destabilize the membrane, contributing to cell instability and neuron death (apoptosis). Both these effects are eventually exerted in a complex frame, with many molecules present: APP N-terminus (before the cleavage); peptides in monomeric, oligomeric and pre-fibrillar assemblies; other cofactors, like metal ions. Thus, even at the monomer level, the interactions between amyloid peptides and biological membranes are still poorly understood [7]. More complete models are required to contribute to recent views of APP and A*β*, where A*β* aggregation is interpreted as a loss of functional A*β* monomers [8].

Molecular simulations, particularly molecular dynamics (MD), became a standard tool of computational biology to understand molecular interactions in such complex frames [9]. Despite the large number of simulation studies involving A*β* monomers [10, 11], oligomers [12, 13], and fibril-like assemblies [14–18], with all species in contact with membrane models, the role of cofactors abundant in the environment of neurons have seldom been taken into account [19]. Among these cofactors, divalent ions, and especially copper, are relevant for a correct physiology of the synapse [20]. Some of the known facts are summarized below.

1. Copper (Cu) and zinc (Zn) are particularly abundant in the synaptic region, reaching concentrations up to 0.3 mM after functional extracellular release [20–23]. These concentrations are many orders of magnitude larger than that inside the cell, where Cu, for instance, is present in negligible amount as an ion available to interactions [24, 25]. The addressing of APP as a copper mediator has been discovered [26] and lately associated to many neurodegenerative disorders [27, 28].
2. Divalent cations change membrane structure, transport properties [29–31], and reactivity [32], thus possibly promoting protein aggregates resembling ion channels and membrane pores [33, 34].
3. Cu ions in contact with A*β* peptides form catalysts for the production of reactive oxygen species, activating dioxygen molecules [35, 36], and promoting oxidative pathways [37–40].

Because of these important issues, the modeling of interactions of divalent cations with lipid charged and zwitterionic membranes is becoming a challenge [41–43]. Indeed, recent accurate models explain the experimentally observed strong interactions between Ca^2+^ and phosphate groups in POPC bilayers [43].

In this work, we compare, for the first time, models of free and peptide-bound divalent cations in interaction with dimyristoyl-phosphatidylcholine (DMPC) bilayers, with special emphasis on oxidized copper. Polarizable models of interactions between divalent cations and biological macromolecules are still experimental [43]. Even for nucleic acids, the contribution of Mg^2+^ to the stability of tertiary RNA folding is intricate [44]. Overall, it is not trivial starting from an unbound condition to sample bound conditions that are observed experimentally. Copper binding is known to be fluxional and strongly dependent on the environment [45, 46]. Therefore we apply separately two modeling techniques: i) a naive non-bonded model of Mg^2+^ that has been used to model the free energy change for the exchange reaction between the water solution and a protein [47], and for neutralizing RNA phosphate groups [48]; ii) a bonded model of Cu^2+^ that has been applied to describe a well documented binding site for Cu-A*β*(1-42) observed in experiments [49–51] and extensively modelled by MD simulations [36, 52].

The models describe interactions between, respectively, Mg^2+^ aqua-ions, A*β*(1-42), and Cu(II)-A*β*(1-42) monomers with DMPC bilayers, the latter a well studied molecular model of biological membrane. The simple model used for the Mg divalent aqua-ion [47, 48] can depict a first approximation of the effects of Cu^2+^ ions, that have size similar to Mg^2+^ when not bound to proteins. These effects mimic those of oxidized Cu on the membrane structure when Cu is released in the synapse and not captured by Cu extracellular carriers.

The model, investigated by means of multiple conventional MD simulations (CMD, hereafter) and replica exchange MD (REMD), is limited to A*β* monomers and to exogenous addition of A*β* to the lipid membrane, rather than to peptide incorporation into membrane during its assembly (see Methods). This assumption is representative of the functional conditions in the synaptic region, where both A*β* peptides and divalent cations are released, at low concentration, in the synaptic cleft during the normal activity of neurons. Also, *in vitro* experiments about A*β*-DMPC interactions mediated by divalent cations have been performed mimicking exogenous addition [53, 54].

Finally, the role of divalent cations in cell signaling is more general than in synapse [55]. Therefore, it is of utmost importance to understand interactions of divalent cations with neuron membrane in the presence of modulating ions’ ligands.

## Methods

A summary of the simulations performed in this work is reported in Table 1.

**Table 1.**
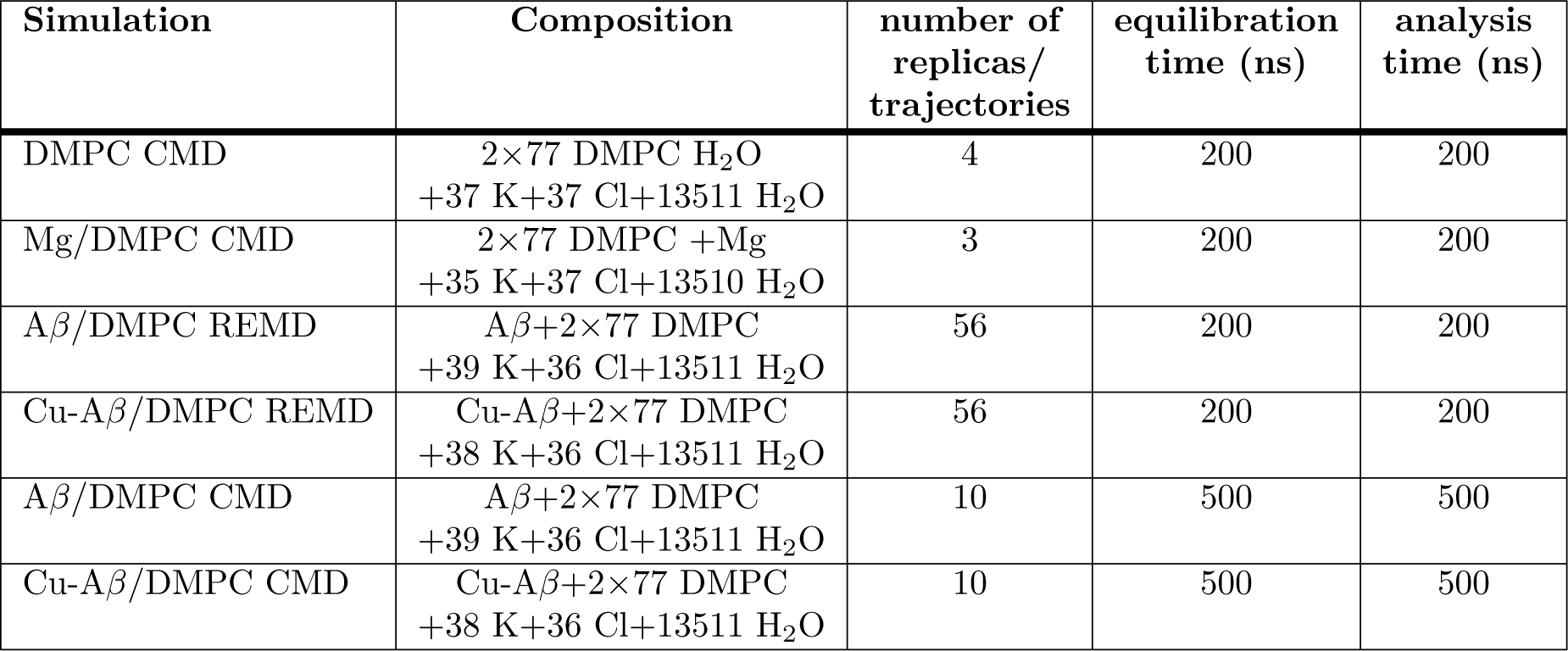
Summary of simulations analyzed in this work. Abbreviations: CMD - conventional molecular dynamics; REMD - replica exchange molecular dynamics; DMPC - dimyristoyl-phosphatidylcholine; A*β* - A*β*(1-42) peptide, charge −3; Cu-A*β* - Cu-A*β*(1-42) complex, charge −2. See Methods for details.

### Set-up of molecular dynamics simulations

The amyloid-*β* peptide of 42 residues (A*β*(1-42)), with and without a single bound copper ion in 2+ oxidation state (Cu^2+^), was simulated with constant temperature molecular dynamics (conventional MD, CMD hereafter) and with the replica exchange molecular dynamics (REMD) method, in order to sample the configurational statistics at the physiologically relevant temperature of 311 K (38° C). The peptide and the ions were put in contact with a bilayer composed of 1,2-dimyristoyl-sn-glycero-3-phosphocholine (also abbreviated as dimyristoyl-phosphatidylcholine, DMPC hereafter) lipid molecules.

The sequence of A*β*(1-42) is:

DAEFRHDSGY_10_ EVHHQKLVFF_20_ AEDVGSDKGA_30_ IIGLMVGGVV_40_ IA with aminoacids indicated with the one-letter code. We used the Amber16 package [56], with the FF14SB [57] force-field for the peptide and monovalent ions (KCl), TIP3P water model [58] for the explicit water solvent, and LIPID14 [59] for the DMPC molecules. AMBER FF14SB force-field is an improved version of FF99SB [60] used in our previous simulations [52, 61]. Although older CHARMM force-fields tend to provide better results for A*β* peptide than old AMBER force-fields [62], new AMBER versions, especially AMBER FF14SB and CHARMM36m provide good agreement with experimental data for A*β* [63, 64]. Moreover, AMBER FF14SB is fully compatible with LIPID14 force-field [59], which is expected to provide optimal accuracy for both lipids and peptide in the simulations. The use of more recent force-fields for IDPs will be pursued in the future, after a detailed comparison between experiments and simulations in generalized ensembles will be reported in the context of amyloid peptides.

We assumed the physiological (pH∼7) protonation state for aminoacid sidechains and free termini. Thus, the charge of A*β*(1-42) is −3 (the N-terminus is protonated and the C-terminus deprotonated). The parameters for copper and copper-bound aminoacids were the same used in our previous MD and REMD simulations [52, 61]. Cu is bound to N and O of Asp 1, N*δ* of His 6 and N_*∊*_ of His 13, the latter protonated at N*δ*. His 14 is neutral and protonated in N_*∊*_, like His 6.

Bond distances and angles involving Cu contribute to harmonic energy terms, with stretching constants, bending constants, and equilibrium values set as fitting parameters of quantum-mechanics calculations at the density-functional level of approximation for truncated models (see Methods in [52]). All the dihedral angles where Cu has index 2 or 3, do not contribute to the potential energy, while those with Cu with index 1 or 4 are obtained by the AMBER99SB force-field where heavy atoms have the same dihedral indices of Cu. Point charges are derived from the restrained electrostatic potential (RESP) method [65, 66], where the electrostatic potential mapped onto the solvent-accessible surface was obtained at the density-functional level of truncated models (see Ref. 52 for details). Excess of net charge, obtained by merging point charges of truncated models into AMBER FF14SB aminoacids, was distributed to C*β* and H*β* of Asp 1, His 6 and His 13 when these residues are bound to Cu^2+^. Lennard-Jones parameters for Cu are reported in the literature [67]. The Cu^2+^ coordination geometry in this empirical force-field is approximately square-planar, with a fifth axial coordination always available to electrostatic interactions, as shown in previous simulations performed with the same force-field [36]. The distance between configurations obtained with this empirical force-field and minimal-energy configurations obtained including explicit electrons (like in density-functional theory applied to truncated models) is small.

As for the free divalent cation, we used the so called “dummy” cation model for Mg^2+^ [47]. This model has been used together with AMBER99SB phosphate groups [48], where it showed reasonable electrostatic properties. Even though this model is a very crude approximation of divalent cations, it is far more reliable than a single site with point charge 2+. A comparison between the affinity of divalent and monovalent cations for the DMPC membrane has been performed by umbrella sampling estimates of free energy differences (see Supporting Information, file).

An initial lattice model of DMPC bilayer was built, using 77 DMPC molecules per layer, with an approximate area per molecule of 62 Å^2^. An orthorhombic simulation cell was built, with the cell side along zeta, the latter the direction normal to the DMPC layer, initially set to 70 Å. The space between the periodic images of the bilayer was filled with 13511 water molecules, initially at the density of 1 g/cm^3^, according to the TIP3P model of bulk water at room conditions. KCl was added in the same space, according to an approximate bulk concentration of 0.1 M. Ions were added randomly replacing water molecules in the initial configuration. The number of Cl^-^ anions was adapted to the change of net charge due to addition of the peptide (see below). The net charge of the simulation cell was always zero.

Initial configurations of amyloid −*β* monomer, without copper (charge 3-) and with copper (charge 2-, because of N-terminus deprotonation), were inserted in the space filled by the water molecules. The same was done for the single divalent cation. The concentration of divalent cation in this cell is 10 mM, thus in the range of the concentration used for Ca, Mg, Zn and Cu in *in vitro* experiments. With a few exceptions, *in vitro* experiments use concentrations, both of peptide and divalent ions, about 2 orders of magnitude larger than *in vivo* in the synaptic region of CNS neurons (∼10-100*µ*M).

To remove eventual bad contacts produced by each initial configuration set-up, we performed 25000 steps of steepest decent energy minimization, followed by other 25000 steps of conjugate gradient energy minimization.

The initial coordinates for the CMD and REMD simulations are included as Supporting Information in the protein data-bank (PDB) file format (the first configuration) and as compressed (Bzip2) XYZ format. See, ,, ,, and.

### Molecular dynamics simulation protocol

We simulated MD trajectories in the isobaric-isothermal (NPT) statistical ensemble, at the constant temperature of 311 K and at the pressure of 1 Atm. Temperature was controlled by a Langevin thermostat [68] with collision frequency of 2 ps^−1^. Pressure was controlled by a stochastic barostat, with relaxation time of 100 fs. The SHAKE algorithm [69] was applied to constrain bonds involving hydrogen atoms. A cut-off of 10 Å was applied for non-bonded interactions and the particle mesh Ewald algorithm [70] was used to compute long-range Coulomb and van der Waals interactions. The simulation time-step was 2 fs.

In order to increase the sampling, we collected several trajectories for each system, starting from different initial conditions, that is the initial velocities (DMPC) and the position of the solute within the water layer (other systems). Composition of each system and some parameters related to sampling are reported in Table 1.

### Replica-exchange molecular dynamics simulation

The REMD simulation was carried out with 56 replicas (or trajectories) corresponding to 56 temperatures ranging from 273K to 500K. The minimized structure was distributed among 56 replicas, and each replica was equilibrated in 200000 steps at the temperature chosen in the temperature distribution. After equilibration, the replica exchange molecular dynamics simulation started, lasting a total time of 400 ns. The exchange of temperature between pair of replicas was attempted every 500 steps of simulation. The REMD simulation is used here mainly to capture the statistical contribution of extended peptide configurations and partially disordered layers, configurations that are rarely sampled at temperatures in the range where the force-field is accurate. The acceptance rate of REMD simulations was, on average, 20% and 21% for, respectively, A*β*(1-42) and Cu-A*β*(1-42).

The behavior of lipid order parameters as a function of temperature (data not shown here) shows that the DMPC bilayer is, at the temperature closest to that of the human body (37° C, 310 K), in the liquid crystalline phase. The configurations sampling the temperature of 311 K are, therefore, analyzed in detail in the following.

To avoid possible bias due to initial configuration construction, we used approximately the second half of each simulation for analysis (see Table 1).

### Analysis

#### Structural properties

Root-mean-square deviation (RMSD) and radius of gyration (*R*_*g*_) were calculated for all A*β*(1-42) atoms using the initial A*β*(1-42) structure as a reference for the RMSD measurement. Secondary structure of A*β*(1-42) was analyzed using DSSP software included in the cpptraj tool [56], a part of AmberTools package. Three regular types of secondary structure were distinguished in the analysis: helices (*α*, 3-10, and *π*), *β*-sheets (parallel and antiparallel), and turns, while the residues in other conformations were treated as unstructured (coil). Solvent accessible surface area was calculated for A*β*(1-42) and lipids using Linear Combinations of terms composed from Pairwise Overlaps (LCPO) method [71], implemented in cpptraj.

Radial distribution function (RDF) measures the probability to have the distance between two sites within a given distance range, *N* (*r*). As usual for liquids and polymers, this quantity is then divided for the same probability for the ideal gas with the same uniform density of sites, *N*_*id*_(*r*): *g*(*r*) = *N* (*r*)*/N*_*id*_(*r*). The function *g*(*r*) approaches the limit *g*(*r*) =1 when *r* → ∞, *i*.*e*. when the two sites in the pair become not correlated.

The bilayer thickness is defined as the distance between the two planes formed by phosphor atoms belonging to each layer. The roughness of a layer is defined as the standard deviation of *z* coordinates of phosphor atoms within each layer.

The number of contacts is defined as the count of the usual distance-based step-like variable:

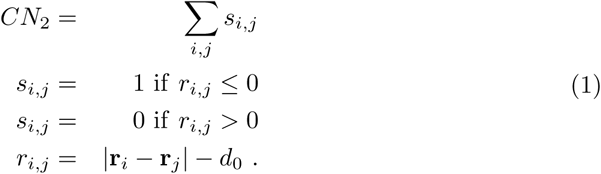

with *i* and *j* running over different sets of atom pairs, each term of the pair contained in a different portion of the system. When the two sets of atoms identify, respectively, atoms belonging to positively charged groups (N*ζ* in Lys and N*η* in Arg) and negatively charged groups (C*γ* in Asp and C*δ* in Glu), we address the contact as an intramolecular salt-bridge. The number of such contacts is indicated as SB and the *d*_0_ parameter is chosen as 4 Å. As for generic inter-residue contacts, we measured the distance between the centers of mass of sidechains in the two involved residues. In this case, *d*_0_ is chosen as 6.5 Å. When the contact between aminoacids and lipid molecules is addressed, the center of mass of DMPC molecules is used and the *d*_0_ distance is 4.5 Å.

### Elastic moduli

Elastic moduli of lipid bilayer were calculated by fitting suitable ensemble averages with the following equations [72]:

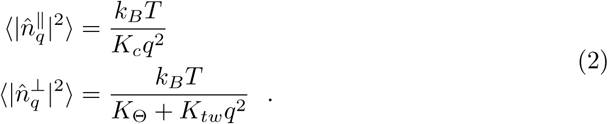

where *K*_*c*_, *K*_Θ_, *K*_*tw*_ are bending, tilt, and twist elastic moduli, respectively, *k*_*B*_ is Boltzmann constant, *T* is temperature, and 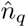 is the reciprocal space vector determined as summarized below (see also supplementary information of Ref. 72 and Ref. 73).

The *xy* plane of the membrane is discretized to a square 8×8 grid. The orientation vector of lipid molecule *j* is 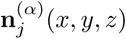 with *α* 1 or 2 for upper and lower layers, respectively. Each vector points from the midpoint between P and C2(glycerol) atoms to the midpoint between the terminal C atoms of the lipid tails. The orientation vectors are projected onto the *xy* plane and are mapped onto the 8×8 grid, providing *n*^(*α*)^(*x, y*). Fast Fourier transform is used to obtain 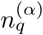, where *q* is the reciprocal space index. From 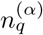 we obtain the quantity

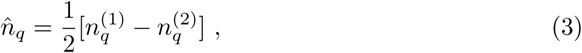

that is decomposed into longitudinal 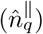 and transverse 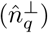 components:

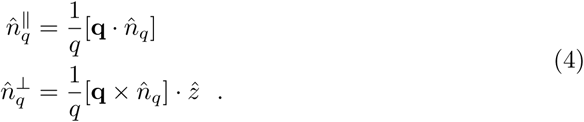

Finally Eq. 2 is used to average according to the collected sampling of lipid molecules.

## Results

### Addition of a divalent cation to the DMPC bilayer

The affinity of Mg^2+^ for the DMPC bilayer was measured by the umbrella sampling method (see Supporting Information, Fig.S1 in). The free energy minimum was found at 17 Å from the bilayer center, thus corresponding to the average distance of P atoms (see below). The flatter shape of free energy around the minimum in the case of Na^+^ is due to the equivalent interactions of Na with phosphate and carbonyl groups of DMPC. These interactions allow a deeper penetration of Na into the bilayer than Mg. The binding free energy of Mg^2+^ was estimated as about four times that of Na^+^. This difference favours the binding of Mg to the DMPC surface compared to Na. This difference is opposite to what expected on the basis of dehydration free energy, that should favour Na compared to Mg, being the hydration free energy at 300 K about five times more negative for Mg compared to Na [74]. This effect is due to the strong electrostatic interactions formed by Mg when absorbed by phosphate groups, together with a significant drift of water molecules towards the bilayer center along with the cation’s penetration. Therefore, interactions with phosphate oxygen and with residual water molecules strongly compensate the loss of water molecules from the Mg first-coordination sphere when Mg is driven from the bulk water towards the bilayer center.

All of the 3 CMD trajectories of Mg/DMPC display a rapid approach of the divalent cation (Mg^2+^) from the bulk to the initially closest layer. After 200 ns, the divalent cation is trapped by phosphate groups of DMPC. Since the 3 CMD trajectories are equivalent in several average properties (like the radial distribution function *g*, see Methods), the average over the 3 trajectories is analyzed in the following. We indicate the cation-bound layer as layer 1 (L1) and the layer not affected by the binding as layer 2 (L2). The difference between *g* calculated for L1 and L2 is displayed in Fig 1. The divalent cation (black) is bound to the phosphate oxygen atoms, thus displaying the coordination distance of 2.9 Å with respect to P atoms. Including the second-shell Patoms (the peak at 3.5 Å), the number of P atoms around the cation is 4. This coordination affects the average distance between charged groups within L1, as it is displayed by the P-P distances (red line), respectively, within each layer L1 and L2. Conversely, atoms farther than P from the perturbing cation are less affected, as shown by the difference in N-P distance distribution among the two layers (blue line).

**Fig 1.**
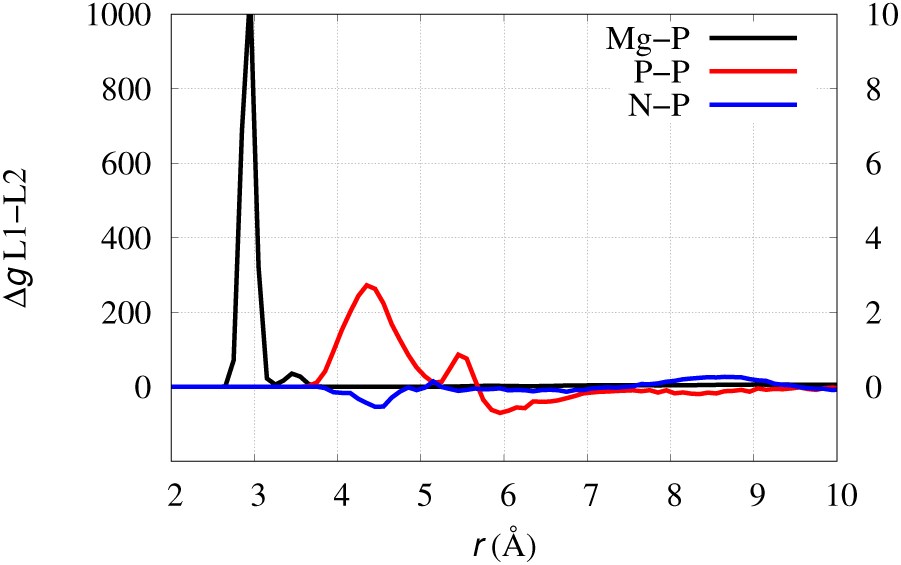
Effect of Mg addition to DMPC. Difference between radial distribution function (*g*) computed in Mg/DMPC for layer 1 (Mg-bound) and layer 2. Mg-P (black line); P-P (red line); N-P (blue line). Left *y*-axis is for black line, right *y*-axis is for red and blue lines.

The formation of a cluster of phosphate groups in L1 induces the release of the electrostatic interactions within the head groups in each layer. Therefore, a consequence of the phosphate neutralization by Mg binding to L1, is a change in the distribution of monovalent counterions at the interface of the two different layers. This effect is emphasized by plotting the difference in K-P radial distribution function between the two layers and by comparing this quantity with the same quantity computed in the absence of divalent cation. In Fig 2A it can be noticed that the distribution of K^+^ in the presence of Mg (black curves) is more asymmetric than with no Mg (red curves). The low symmetry of K-P distribution in the absence of Mg (red curves) is due to sampling limitations. Indeed, the presence of Mg on the L1 layer displays a “hole” in K distribution where there is a little excess in the absence of Mg. Because of the change in interactions between K^+^ and P at short distance (the peaks at the left), there is also a decrease of bulk concentration within a distance of 1 nm from the P atoms. This change of the electrostatic properties between the two sides of the bilayer is equivalent to a weak polarization of the membrane. This asymmetry is caused by the asymmetry in the P-P radial distribution (Fig 2B), that is due to the formation of the Mg-O(P) coordination.

**Fig 2.**
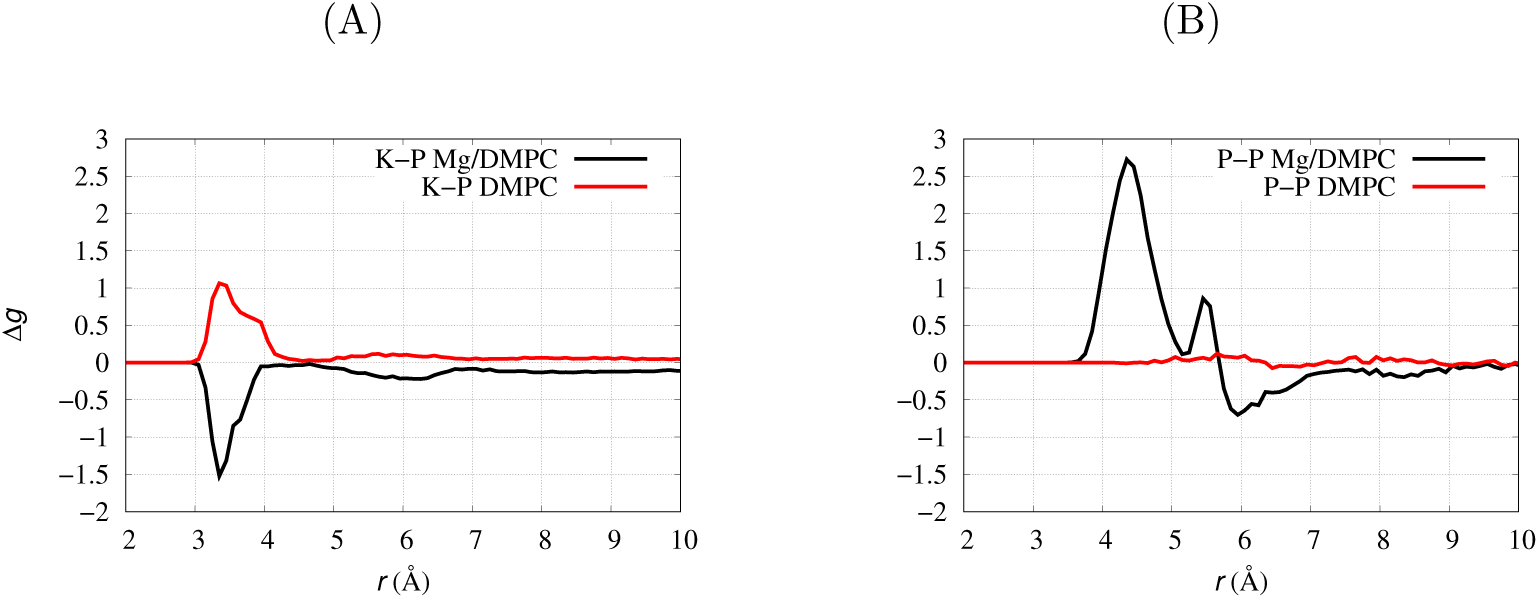
Effect of Mg addition to DMPC. Difference between radial distribution function (Δ*g*) computed for layer 1 (Mg-bound) and layer 2. (A) K-P in Mg/DMPC (black line, average of trajectories 1-3); K-P in DMPC (red line, trajectories 1-4). (B) P-P in Mg/DMPC (black line, trajectories 1-3); P-P in DMPC (red line, trajectories 1-4).

The asymmetry of the interactions between divalent cations added from one side of the bilayer is consistent with experimental data reported for exogenous addition of Cu^2+^ and Zn^2+^ to bilayer models (POPC/POPS mixtures) [53]. The comparison between ^2^H and ^31^P ss-NMR spectra of POPC/POPS molecules shows that P atoms are strongly affected, while the molecular tails in the hydrophobic region of the bilayer are almost unaffected. The addition of Cu^2+^ to these membranes induces the formation of smaller vesicles, thus showing a dramatic effect of this ion on the bilayer stability.

The effect of the divalent cation on the elastic property of DMPC is also significant. In Table 2 we report the elastic constants determined by the different simulations, with averages of Eq. 2 (see Methods) computed over all the acquired trajectories (see Table 1).

**Table 2.** Elastic moduli of DMPC bilayer with no addition (DMPC) and interacting with, respectively, a divalent cation (Mg/DMPC), the A*β* peptide (A*β*/DMPC), and the Cu-A*β* peptide (Cu-A*β*/DMPC). Average is computed over 10 windows of 20 ns each, during the last 200 ns of each CMD trajectory. Standard error is within parenthesis.

| Elastic moduli | DMPC | Mg/DMPC | Aβ/DMPC | Cu-Aβ/DMPC |
| --- | --- | --- | --- | --- |
| $K_c$ ($10^{-20}$ J) | 7.859 (0.369) | 14.568 (0.756) | 13.316 (1.307) | 15.210 (2.077) |
| $K_\theta$ ($10^{-20}$ J/nm<sup>2</sup>) | 6.679 (0.191) | 5.200 (0.200) | 6.767 (0.241) | 7.095 (0.200) |
| $K_{tw}$ ($10^{-20}$ J) | 1.447 (0.010) | 1.629 (0.006) | 1.668 (0.061) | 1.668 (0.042) |

The values are in the range of those found in DPPC atomistic simulations [72], though the conditions (temperature, force-field, etc.) are different. The bending constant (*K*_*c*_) of pure DMPC is smaller than in all the other cases, where the DMPC is perturbed by exogenous addition of species. This change shows that the addition of any species on one side of the bilayer increases the rigidity of curvature, because of the change exerted more on one layer than on the opposite layer. On top of this effect, that is due to the asymmetry of the addition, the tilt modulus (*K*_*θ*_) is significantly smaller for Mg/DMPC compared to the DMPC bilayer both unperturbed (DMPC) and with the peptide (A*β*/DMPC and Cu-A*β*/DMPC) floating over the bilayer surface. This additional information reveals that the formation of bridges between phosphate groups occurring in Mg/DMPC (see Fig 1) produces a cluster of 3-4 lipid molecules that changes the elasticity of DMPC. As described above (and also in detail below), the lipid molecules belonging to the cluster are more rigid and create a small hollow in the surface. The perturbation exerted by Mg-phosphate interactions makes a little hollow over the bilayer surface affected by Mg binding. This little hollow can be observed looking at the configurations where the Mg penetration is deep, like in Fig 3. This local perturbation allows the molecules neighbor to the cluster to more easily tilt with respect to the bilayer normal.

**Fig 3.**
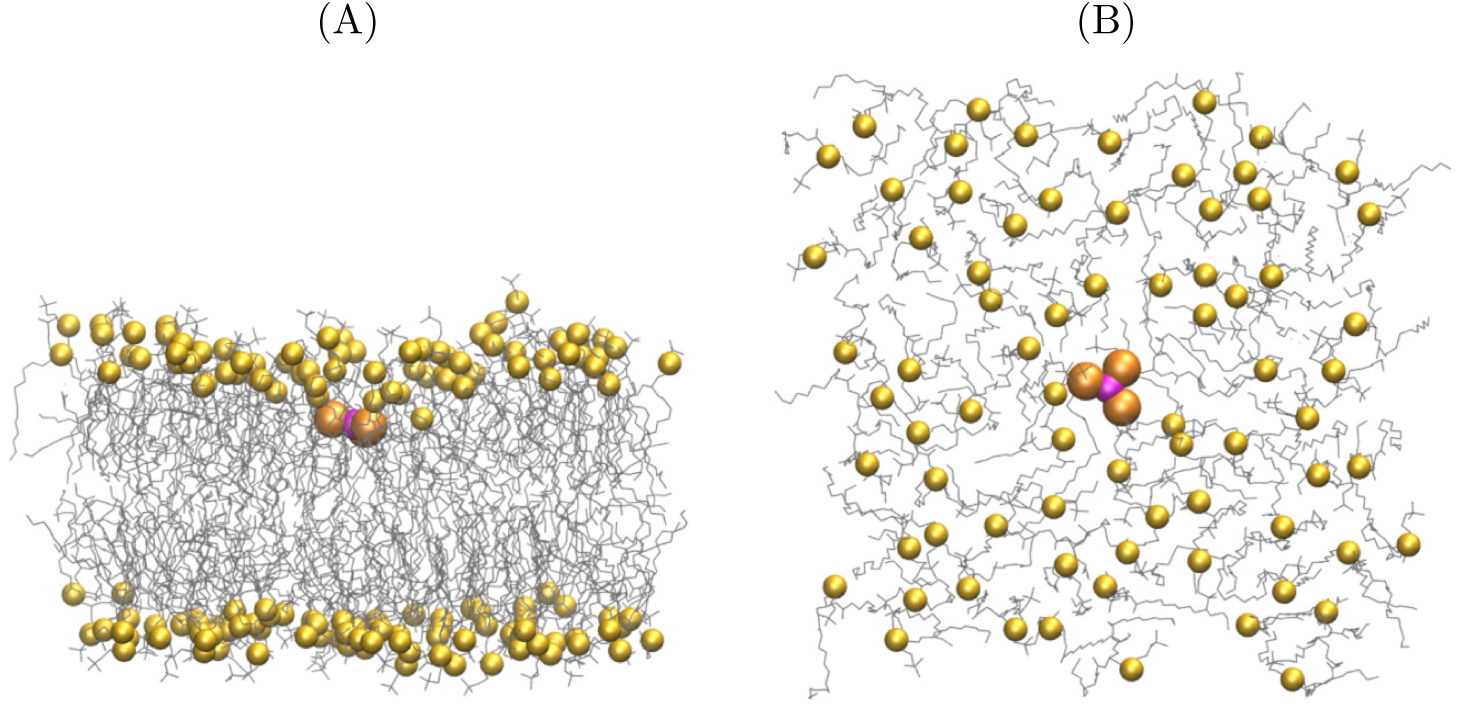
Effect of Mg addition to DMPC. Configuration of Mg/DMPC where the distance between Mg (purple sphere) and the bilayer central plane is minimal along with the CMD simulations 1-3. P atoms in DMPC are represented as yellow spheres, those within 3.5 Å from Mg are emphasized in orange. The other DMPC molecules are represented as thin bonds. Water and KCl are not displayed. Atomic radii are arbitrary. Panel B is the same structure in A observed from the *z* axis and with only lipid molecules in L1 displayed.

The effect of Mg addition to L1 does not significantly alter other structural parameters of the bilayer at the same temperature (see Table 3). For instance, bilayer thickness and area per lipid compare well with the values measured by diffraction studies for DMPC [75]. Experiments report thickness at *T* =303 and 323 K of, respectively, 36.7 and 35.2 Å^2^, while in our MD simulation at 311 K thickness is 34.4 Å^2^. This small difference may be due to the slightly different way used to measure the thickness (see Methods and Ref. 75). The experimental area per lipid is 59.9 and 63.3 Å^2^ at the same two probed temperatures, while it is 63.8 at 311 K. Negligible effects are observed for the average roughness with the Mg^2+^ addition (see Table 3), thus confirming that any effect do to Mg/DMPC association is very localized in space.

**Table 3.** Bilayer structural data averaged over the second half of all trajectories (avg.) and selected trajectories (traj./REMD). Root-mean square errors are within brackets.

| Simulation | Area per lipid (Å<sup>2</sup>) | Thickness (Å) | Roughness L1 (Å) | Roughness L2 (Å) |
| --- | --- | --- | --- | --- |
| DMPC | 63.75(0.05) | 34.4(0.2) | 2.4(0.3) | 2.5(0.4) |
| Mg/DMPC | 63.75(0.05) | 34.3(0.2) | 2.5(0.4) | 2.5(0.4) |
| Aβ/DMPC (REMD) | 64.5(0.1) | 34.4(0.2) | 2.5(0.3) | 2.5(0.3) |
| Cu-Aβ/DMPC (REMD) | 64.4(0.1) | 34.4(0.2) | 2.5(0.3) | 2.5(0.4) |
| Aβ/DMPC (avg.) | 60.6(1.4) | 35.6(0.6) | 2.7(0.5) | 2.7(0.5) |
| Cu-Aβ/DMPC (avg.) | 60.6(1.2) | 35.6(0.6) | 2.7(0.5) | 2.7(0.5) |
| Aβ/DMPC (traj. 1) | 61.6(1.1) | 35.5(0.5) | 2.6(0.4) | 2.6(0.4) |
| Aβ/DMPC (traj. 5) | 60.8(1.6) | 35.4(0.7) | 2.7(0.4) | 2.7(0.5) |
| Cu-Aβ/DMPC (traj. 8) | 60.9(1.1) | 35.6(0.5) | 3.2(0.9) | 3.3(0.9) |

### Exogenous addition of A*β* peptide to the bilayer

In the REMD A*β*/DMPC and Cu-A*β*/DMPC simulations the DMPC bilayer is in the liquid crystal phase at all the probed temperatures, consistently with similar MD simulations reported in the literature [76]. The temperature dependence of the area per lipid in A*β*/DMPC REMD simulation is displayed in Fig 4, together with the available experimental results for DMPC [75], the result for CMD at *T* =311 K for DMPC, and the average of 10 CMD trajectories at *T* = 303 K described below. The behavior for Cu-A*β*/DMPC is not graphically distinct from A*β*/DMPC and, therefore, it is not displayed. The REMD simulation is able to capture the increase of area per lipid (*A*) as *T* increases as well as the area per lipid at high *T*, but it is dominated by high-*T* lipid configurations that are often exchanged in REMD with low-*T* configurations. However, REMD can adequately probe the possibility of peptide penetration at the highest area per lipid accessible, both by experiments and simulations, in the liquid crystal phase of DMPC. Therefore, it is expected that for lower *A* peptide penetration would be more difficult than at high *T*.

**Fig 4.**
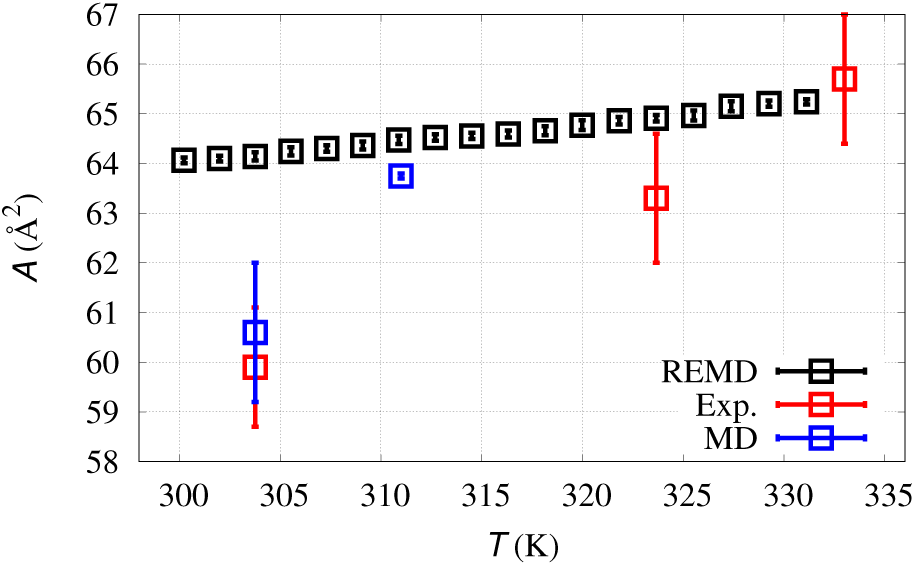
Area per lipid: comparison with experiments. Area per lipid (*A*) as a function of temperature (*T*): average results for REMD simulation (black squares); experimental results at 303, 323, and 333 K (red squares [75]); average of 10 CMD simulations for A*β*/DMPC at 303 K and DMPC at 311 K (blue squares).

In Fig 5 we display the radial distribution function *g* for selected pairs, to show the extent of penetration of N- and C-termini (respectively Nt and Ct) through the membrane surface (using P atoms in the pair) or towards the membrane center (using the terminal C atom in the two acyl chains of DMPC, Cf hereafter). The *g* function is measured at *T* =311 K, *i*.*e*. the physiological temperature of the synaptic membrane. The REMD trajectory at 311 K shows that the propensity for A*β* and Cu-A*β* N-termini to interact with the membrane surface is limited to the head groups of the DMPC bilayer, the P atoms. The peaks in Fig 5A (black lines for A*β*/DMPC) represent the electrostatic interaction between the positively charged Nt group of A*β* with the negatively charged phosphate groups (see also the number of salt-bridges discussed below). The peptide N-terminus (residues 1-16) contains most of the charged sidechains and it is the peptide segment involved in metal ion binding. For this reason the behavior of N- and C-termini are expected to be different when they are in contact with a charged membrane. The approximate symmetry of the *g* function measured for different layers in the bilayer membrane (L1 and L2) shows that in both conditions the N-terminus of the peptide is floating above the membrane surface, going back and forth from one layer to the other. The lower symmetry of A*β*/DMPC (black lines) compared to Cu-A*β*/DMPC (red lines) shows that even the wide REMD sampling is not fully adequate to capture the intrinsic symmetry of the system when electrostatic interactions occur.

**Fig 5.**
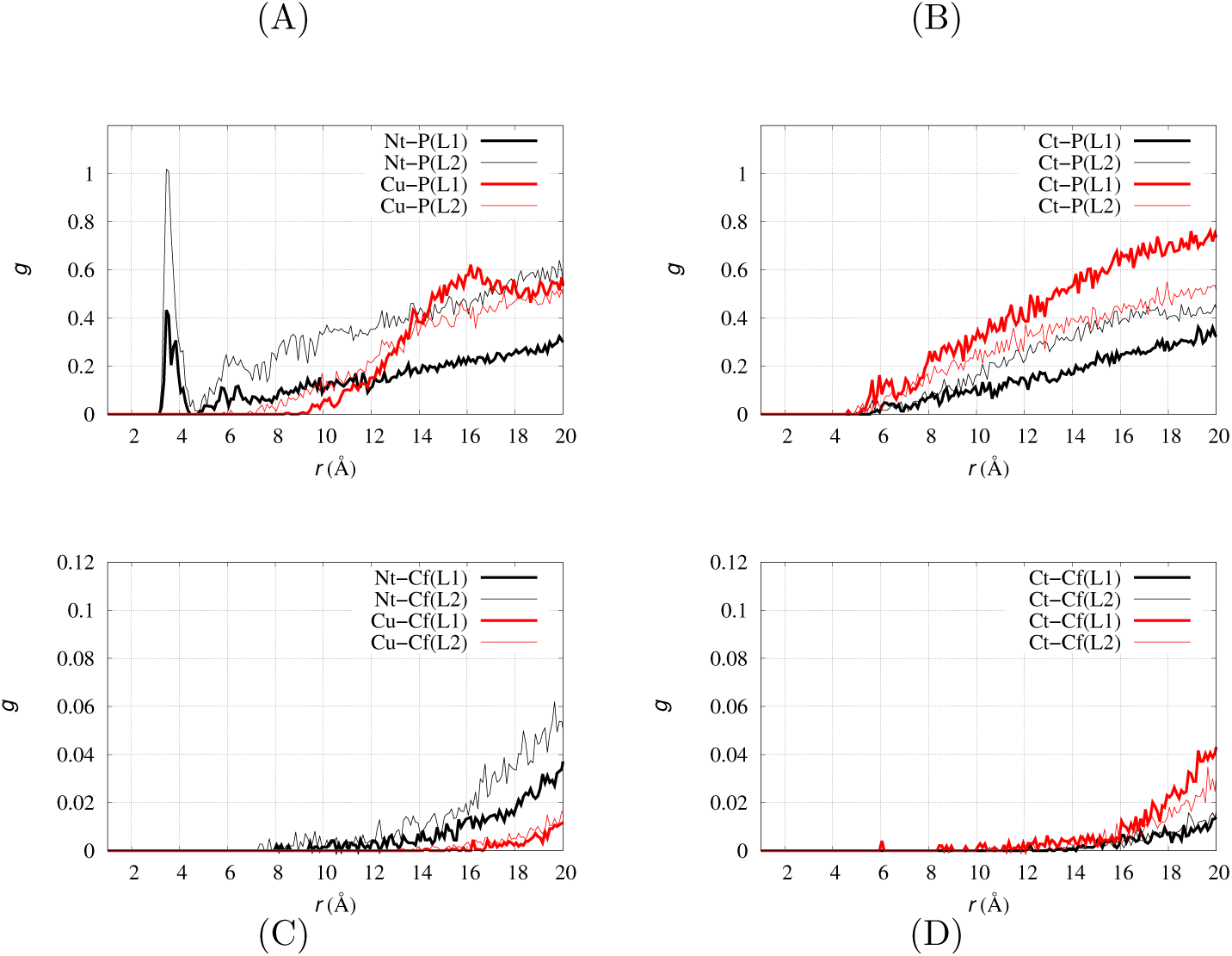
Radial distribution functions by REMD. Radial distribution function (*g*) computed for layer 1 (thick line) and layer 2 (thin line) in REMD simulations for configurations at *T* =311 K. Notice that *y* range in bottom panels is 1/10 of that in top panels. (A) Nt-P in A*β*/DMPC (black lines) and Cu-P in Cu-A*β*/DMPC (red lines); (B) Ct-P in A*β*/DMPC (black lines) and Ct-P in Cu-A*β*/DMPC (red lines); (C) Nt-Cf in A*β*/DMPC (black lines) and Cu-Cf in Cu-A*β*/DMPC (red lines); (D) Ct-Cf in A*β*/DMPC (black lines) and Ct-Cf in Cu-A*β*/DMPC (red lines).

The A*β* peptide Nt atom approaches the P atoms at 3.5 Å, while Cu in Cu-A*β* rarely reaches a distance lower than 6.5 Å. The Cu-binding to A*β* reduces the interactions between the N-terminal region of the A*β* peptide and DMPC head groups, producing a more symmetric *g* function among the two layers. This effect is expected since the interaction with Cu spreads the positive charge over the Cu-bound residues, while in the charged N-terminus (when not bound to Cu) of the A*β* peptide, the positive charge density is higher and the interactions with negatively charged groups at the bilayer interface are more likely.

The peptide rarely penetrates the membrane bilayer, as shown by the *g* function for pairs involving the Cf atoms (the bottom of the acyl chains in lipid molecules, Figs 5C-D). According to bilayer structure (see results reported below) the average distance between P atoms and the center of the bilayer is about 17 Å. Therefore, the Nt atom for A*β*/DMPC (black lines in panel C) and the Ct atom in Cu-A*β*/DMPC (red lines in panel D) significantly approach the bilayer center, showing a deep penetration in rare configurations in the trajectory. Noticeably, when Cu is bound to the peptide (red lines) penetration occurs from the C-terminus, while when Cu is absent the N-terminus is allowed to move from the surface (P atoms) towards the bilayer center. The representation of this change in penetration is better understood examining the few snapshots contributing to the contribution to *g* at short distances in, respectively, Cf-Nt (A*β*/DMPC, Fig 5C) and Cf-Ct (Cu-A*β*/DMPC, Fig 5D). In Fig 6 we display, left and right panels, one of such configurations for, respectively, each of the two systems. It can be observed that a common feature of the peptide structure in these configurations is the breaking of cross-talk between the N- and C-termini. This cross-talk is always present when the peptide (both A*β* and Cu-A*β*) are in water solution and it is often maintained when the peptide interacts with the membrane surface. The interplay between the release of intra-peptide interactions and penetration into the bilayer is discussed in more detail below.

**Fig 6.**
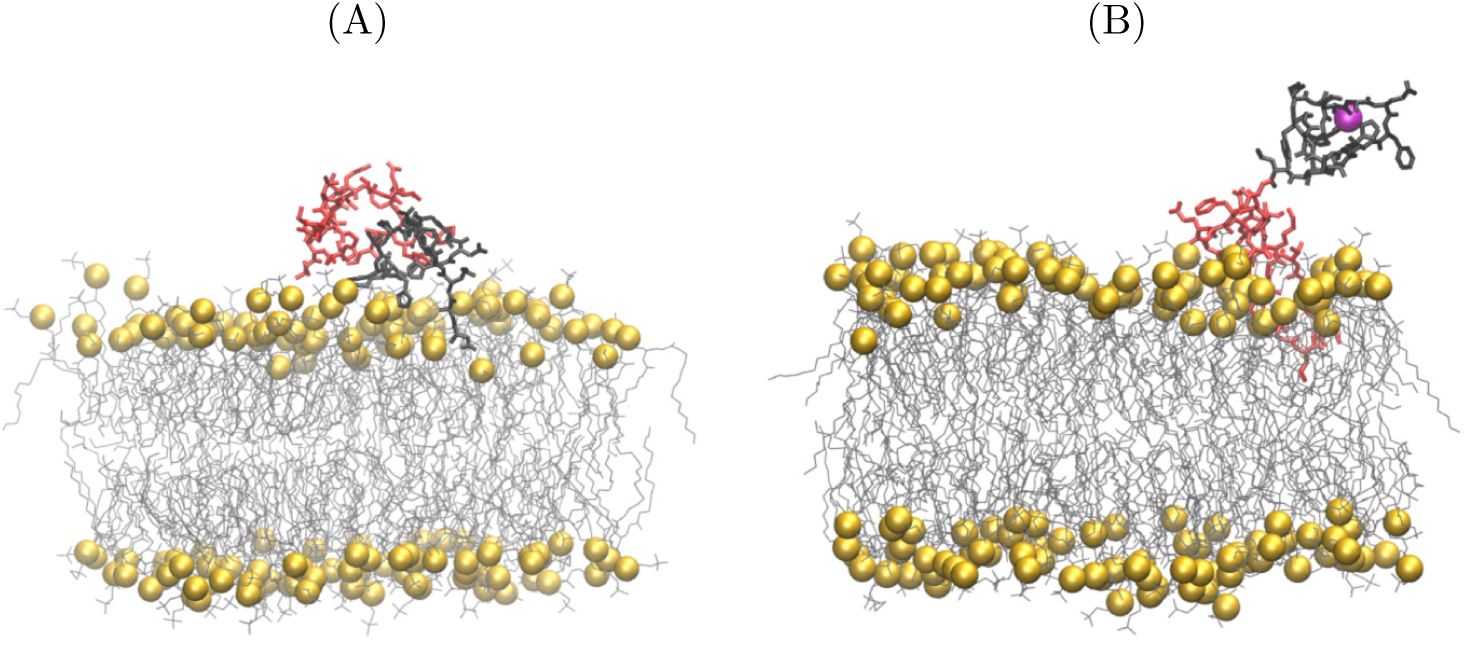
Penetration into DMPC. Configurations of A*β*/DMPC (left) and Cu-A*β*/DMPC (right) displaying the deepest penetration into the lipid bilayer in REMD simulations. The configurations are those where the distance between any peptide atom and any of the bilayer Cf atoms (the terminal methyl group of acyl DMPC sidechains) is minimal along with the trajectory at *T* =311 K. The peptide is represented as bonds (N-terminal residues 1-16 in black, C-terminal residues 17-42 in red), Cu as a purple sphere. P atoms in DMPC are represented as yellow spheres. The other DMPC molecules are represented as thin bonds. Water and KCl are not displayed. Atomic and bond radii are arbitrary.

The number of intramolecular salt-bridges (SB) within the peptide (Table 4) is consistent with the data reported for the simulation of the same peptides in water (last columns). For A*β*/DMPC SB is similar to the value in water, with N(Asp 1) providing a contribution of approximately 1 in both cases. This shows that despite the few interactions between the N terminus and the phosphate groups of DMPC, the intramolecular salt-bridge involving N(Asp 1) in the peptide is not statistically broken and the monomeric peptide keeps the network of intramolecular salt-bridges almost intact. This result is consistent with the rare events of membrane penetration observed in REMD at *T* =311 K. Also in Cu-A*β*/DMPC SB does not change with respect to the value in water. These data show that the N-terminus of A*β*(1-42) and Cu-A*β*(1-42) is bent towards the peptide by, respectively, intramolecular salt-bridges and covalent bonds involving Cu. Thus, N-terminus is rarely released by the peptide cross-talk to form new interactions with the DMPC phosphate groups.

**Table 4.**
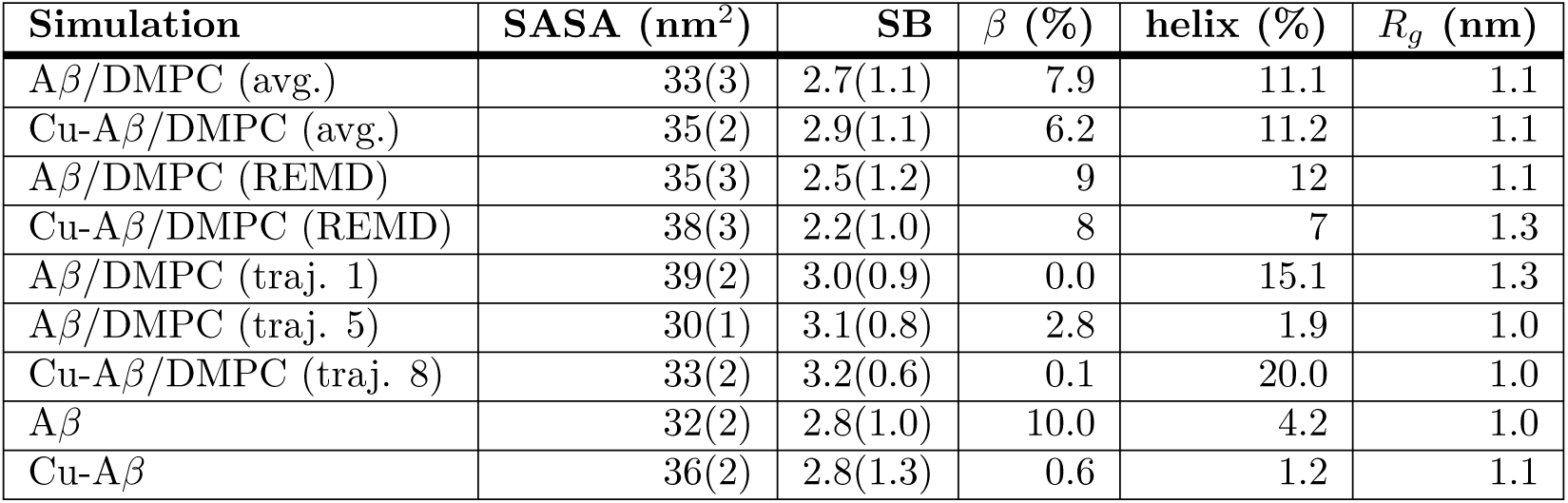
Structural data averaged over the second half of all trajectories (avg.) and selected trajectories (traj./REMD). See Methods for definitions. Root-mean square errors are withing brackets.

| Simulation | SASA (nm<sup>2</sup>) | SB | $\beta$ (%) | helix (%) | $R_g$ (nm) |
| --- | --- | --- | --- | --- | --- |
| Aβ/DMPC (avg.) | 33(3) | 2.7(1.1) | 7.9 | 11.1 | 1.1 |
| Cu-Aβ/DMPC (avg.) | 35(2) | 2.9(1.1) | 6.2 | 11.2 | 1.1 |
| Aβ/DMPC (REMD) | 35(3) | 2.5(1.2) | 9 | 12 | 1.1 |
| Cu-Aβ/DMPC (REMD) | 38(3) | 2.2(1.0) | 8 | 7 | 1.3 |
| Aβ/DMPC (traj. 1) | 39(2) | 3.0(0.9) | 0.0 | 15.1 | 1.3 |
| Aβ/DMPC (traj. 5) | 30(1) | 3.1(0.8) | 2.8 | 1.9 | 1.0 |
| Cu-Aβ/DMPC (traj. 8) | 33(2) | 3.2(0.6) | 0.1 | 20.0 | 1.0 |
| Aβ | 32(2) | 2.8(1.0) | 10.0 | 4.2 | 1.0 |
| Cu-Aβ | 36(2) | 2.8(1.3) | 0.6 | 1.2 | 1.1 |

The bilayer structure (Table 3) shows only a moderate propensity for larger thermal fluctuations, induced by the perturbation due to weak interactions with the peptide, and a small increase in thickness.

Because of the extended conformational sampling in REMD, in both cases the peptide N-terminus moves back and forth between the two layers, because of the usual periodic boundary conditions used in simulations. As a consequence of the weak interactions between the peptide and the DMPC bilayer, the distributions of K-P and P-P distances are approximately symmetric among the two layers and almost identical to those of pure DMPC (data not shown here). The peptide does not change the distribution of monovalent ions.

In order to extract more information about possible specific interactions favoring asymmetry in structural and electrostatic properties among the two layers, in the following we compare 10 separated long (1 *µ*s) CMD simulations performed for both A*β*/DMPC and Cu-A*β*/DMPC models.

### Comparing different peptide/DMPC associations

In this section, the NPT-ensemble MD simulations (that we indicate as conventional MD, CMD) of A*β*/DMPC and Cu-A*β*/DMPC are described. Since the sampling in CMD is more limited than in REMD, the different trajectories allow a comparison between different kinds of A*β*/DMPC and Cu-A*β*/DMPC association.

In Fig 7, in order to describe the type of association, the distance along the *z* axis between the bilayer center and the closest atom of the peptide is displayed as a function of time for all trajectories. Among 10 1 *µ*s-long trajectories acquired for each of the two species, A*β*/DMPC (panel A) and Cu-A*β*/DMPC (panel B), respectively, we observe the rapid incorporation of the peptide into the bilayer in one trajectory only, trajectory 1 of A*β*/DMPC. As for A*β*/DMPC, we observe a partial incorporation after 600 ns for trajectory 5, while for Cu-A*β*/DMPC a moderate bilayer penetration is observed for trajectory 8. These data show that in most of the cases the peptide interacts with head groups (around P atoms). On average, the distance between Cu and the center of the membrane is 42.0±10.6 Å for Cu-A*β*/DMPC compared to 15.3±2.4 Å for Mg in Mg/DMPC. In all simulations the bilayer thickness is about 34 Å (see Table 3 and discussion below), thus the average distance between P atoms and the central plane of the bilayers is never below 17 Å. The approach of Mg towards the bilayer central plane does not significantly drift, on average, the P atoms towards the center of the bilayer, because the density of P atoms projected along the *z* axis does not change (data not shown here). However, as described above, the perturbation makes a little hollow over the bilayer surface affected by Mg binding (see Fig 3 and discussion above).

**Fig 7.**
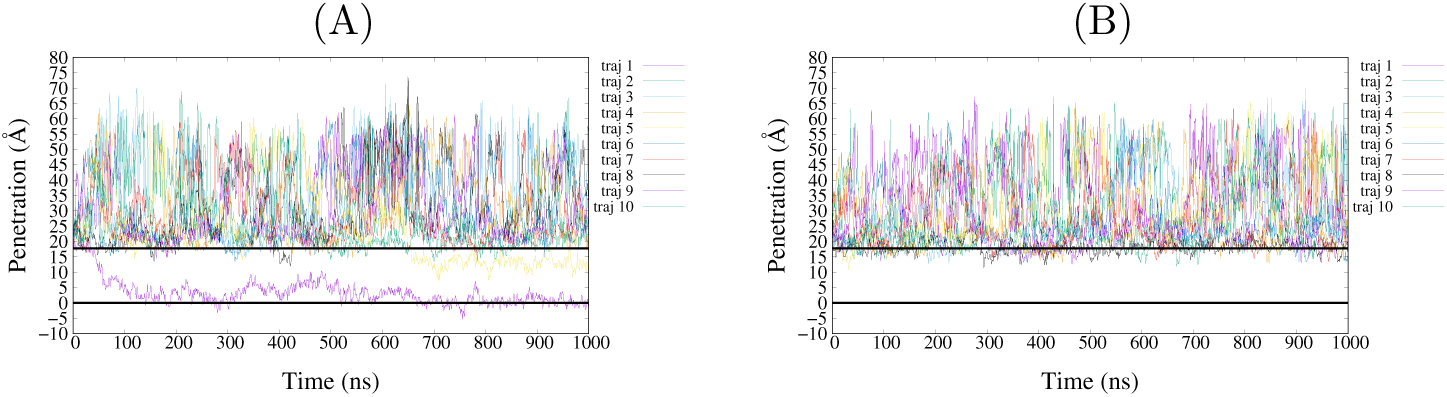
Penetration into DMPC. Penetration of A*β*(1-42) (left, A*β*/DMPC) and Cu-A*β*(1-42) (right, Cu-A*β*/DMPC) into the lipid bilayer. *y* axis is the *z* coordinate of the lowest atom (minimal *z*) of peptide. The horizontal line at *y*=0 indicates the center of geometry of the bilayer which is the average of *z* coordinates of all DMPC’s atoms. The horizontal line at 17.7 Å shows the average position of all P atoms.

These observations are consistent with the experimental data reported for exogenous addition of A*β*(1-42) to bilayer models (POPC/POPS mixtures) [53]. Comparing ^2^H and ^31^P solid state-NMR of A*β*(1-42) and Cu-A*β*(1-42), a neat indication of the confinement of peptides around the head groups is shown. Peptide incorporation during the bilayer preparation, on the other hand, has more severe impact on NMR data and bilayer stability, irrespective of Cu addition.

### Effect of peptide addition to DMPC bilayer structure

The area per lipid as a function of temperature measured by REMD simulation (see above) and consistent with experimental data [75] shows that the area per lipid increases with temperature. Therefore, most of the changes displayed in Table 3 are due to the lower *T* used in the CMD simulations of A*β*/DMPC and Cu-A*β*/DMPC (*T* =303 compared to DMPC and Mg/DMPC (*T* =311 K). The choice of *T* =303 K is to compare these results to CMD simulations of A*β*(1-42) and Cu-A*β*(1-42) in the absence of DMPC [52]. Despite the more significant effect of peptide/DMPC interactions in the 10 separated CMD than in REMD, the changes in bilayer structural parameters (Table 3) are consistent with the experimental data [53] that show a small structural effect for the bilayer, when addition of both A*β*(1-42) and Cu-A*β*(1-42) to the POPC/POPS bilayer is exogenous. On the other hand, the peptide incorporation has a more significant effect on the structure of DMPC head groups, as it discussed in the next subsections. As for bilayer thickness, in our simulations we observe a few incorporated samples, but in all cases where peptide incorporation occurs the thickness of the bilayer is not dramatically affected, compared to the case where the peptide is confined at the membrane surface. The change in area per lipid is, on the other hand, more significant for trajectory 1 (61.6 Å^2^) compared to the average (60.6). This shows that peptide digs a little hollow separating the lipid molecules one from each other, with no wide changes in the bilayer structure, like those emerging from the displacement of a lipid head group from the layer to the solvent.

The order parameters of hydrophobic DMPC chains (data not shown here) show a negligible effect of both A*β* and Cu-A*β* exogenous addition to DMPC. This is an expected effect, since the penetration of the peptide into the bilayer is small (see Fig 7B).

### Effect of peptide addition to DMPC on electrostatic properties

We extend the measure of the effects of interactions between peptide and the DMPC head groups on the distribution of monovalent ions (K^+^) on the two layers. Again, to better understand these effects we analyze the different CMD trajectories. In Fig 8, we compare the radial distribution function for pairs involving P atoms in DMPC and atoms in the N-terminus of the peptide, N(Asp 1) and Cu in, respectively, A*β*/DMPC and Cu-A*β*/DMPC. For instance, comparing trajectories 1 and 2 for A*β*/DMPC and Cu-A*β*/DMPC, we notice that the more symmetric is the interaction between the peptide among the two layers (left panels), the more symmetric is the distribution of K^+^ (right panels). It is also interesting to notice that the strong interaction of trajectory 1 for A*β*/DMPC (see above), produces a polarization of K^+^ that is opposite to that produced by Mg^2+^ (Fig 2A, black curve).

**Fig 8.**
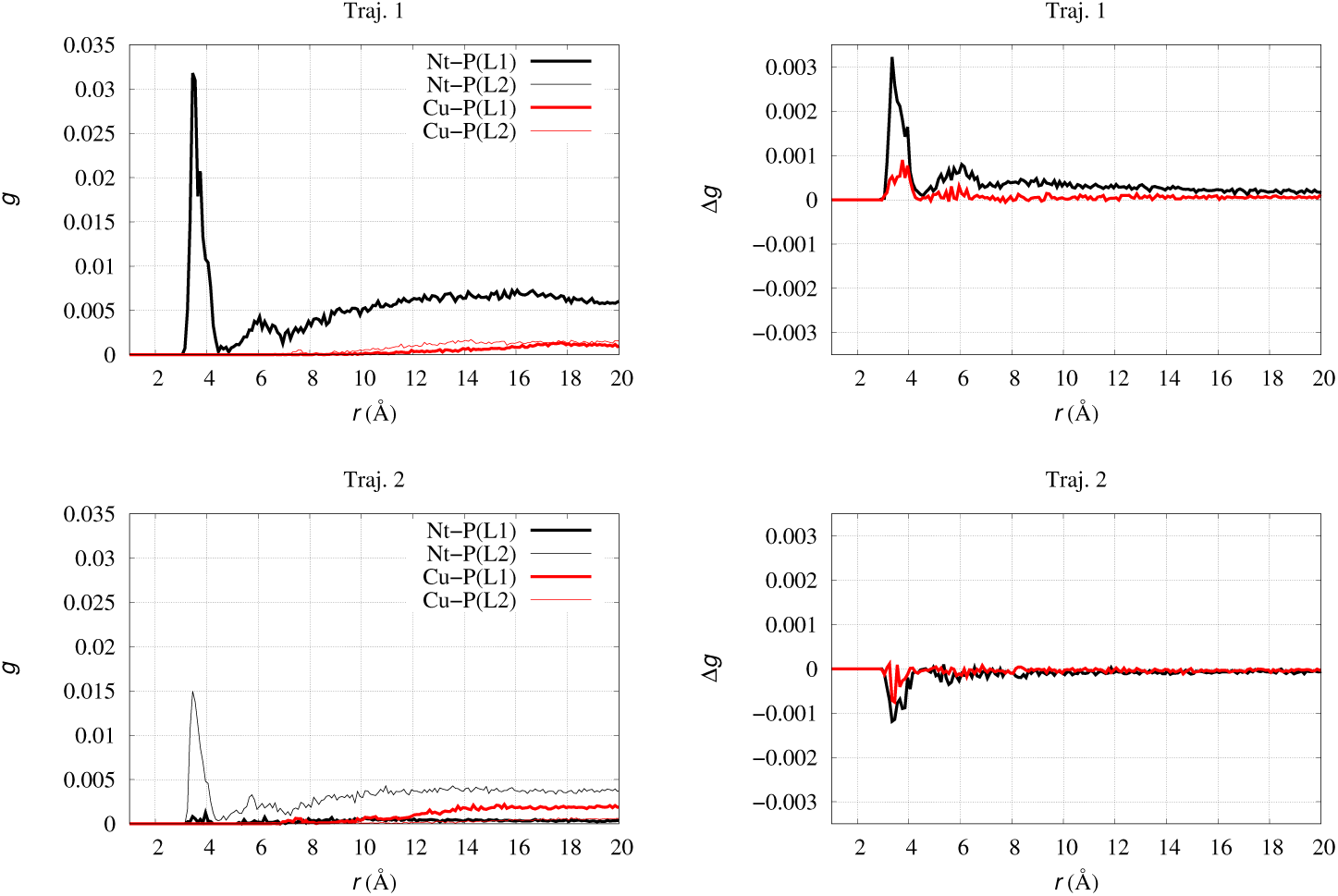
Radial distribution function. Radial distribution function (*g*, left panels) and radial distribution difference (Δ*g*, right panels) computed for layer 1 (thick line) and layer 2 (thin line). Nt-P in A*β*/DMPC (black line, left); Cu-P in Cu-A*β*/DMPC (red line, left); K-P in A*β*/DMPC (black line, right); K-P in Cu-A*β*/DMPC (red line, right); Trajectory 1 (top); trajectory 2 (bottom).

### Effect of Cu and DMPC on peptide structure

Circular dichroism provides important experimental information about the change of structure of A*β*(1-42) and Cu-A*β*(1-42) when the peptides are added to the preformed bilayer [53]. When these experiments are performed at low peptide concentration (by using synchrotron radiation sources), aggregation phenomena are minimized during the measurements. These experiments show that the change of structure of the peptide is minimal, both without and with Cu, when peptides are added to the bilayer. A more significant change occurs when peptides are incorporated during bilayer formation and, in the latter case, the addition of Cu is also affecting the structural modification. In Fig 9 we report the average secondary structure of the peptide, both without DMPC (top, data from Ref. 52) and with DMPC (bottom, this work). The data show that the effect of DMPC association on the peptide is on average small: there is only a significant increase in population of helical regions together with a spreading of the *β*-sheet content among residues. We notice that simulations with no membrane have been performed with a different force-field (AMBER FF99SB).

**Fig 9.**
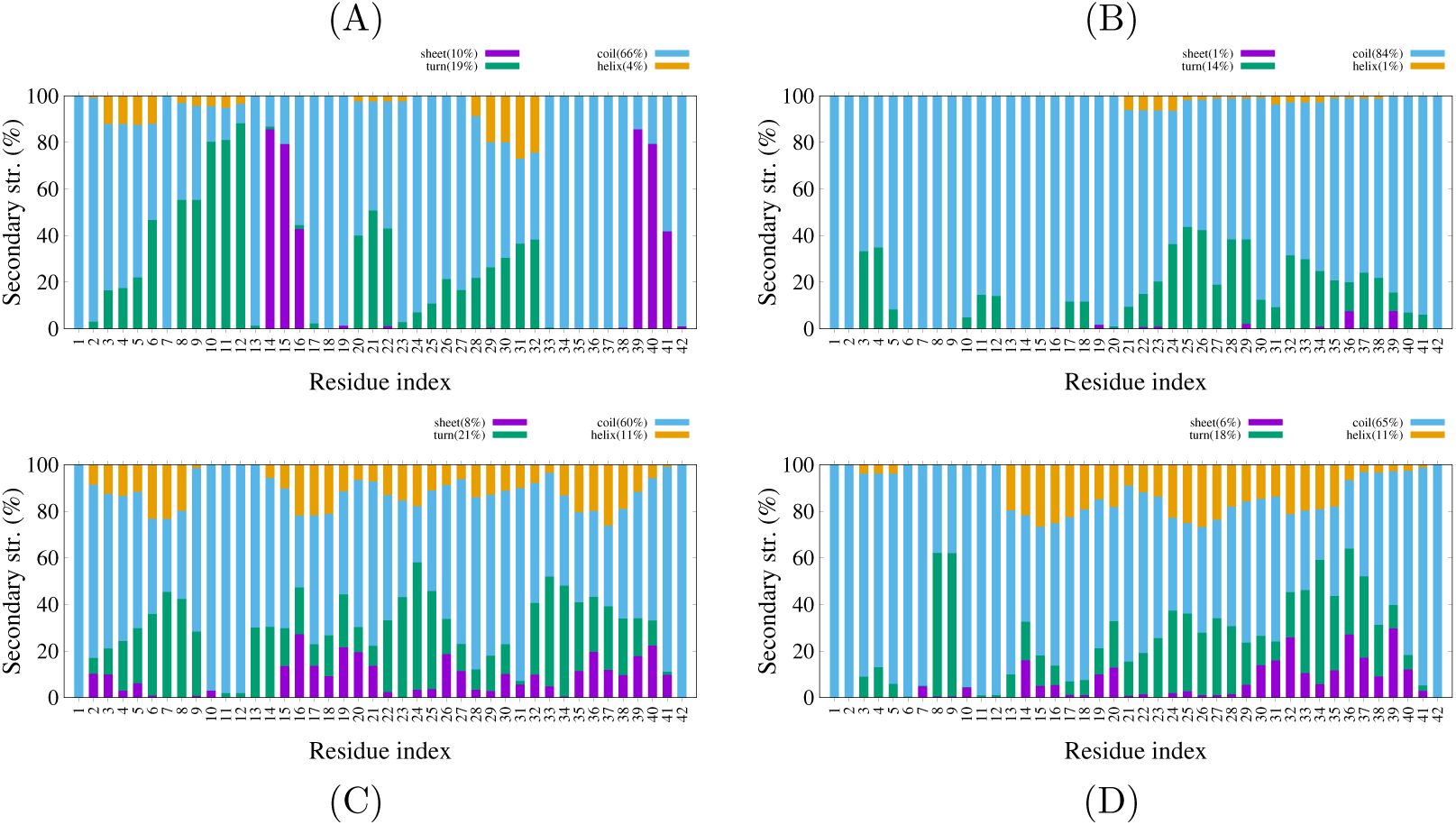
Secondary structure. Secondary structure (see Methods for definition) as a function of residue in A*β*. Top: (A) A*β*(1-42) and Cu-A*β*(1-42) (B) without DMPC [52]. Bottom: secondary structure averaged over 10 trajectories, A*β*(1-42)/DMPC (C) and Cu-A*β*(1-42)/DMPC (D).

In Table 4 we compare structural parameters averaged over 10 trajectories, with those obtained for some selected trajectories, the latter showing the largest extent of association with DMPC. As for those trajectories that are more strongly interacting with the bilayer (especially trajectories 1 of A*β*/DMPC and 8 of Cu-A*β*/DMPC) the helical content is significantly increased. This is an expected result, since it is well known that the incorporation of A*β*(1-40) into vesicles produces *α*-helical motifs in the peptide [77]. It must be noticed that when the peptide is embedded into the bilayer (A*β*/DMPC, traj. 1) there is an expansion of the peptide, while the association with the bilayer surface (A*β*/DMPC, traj. 5, Cu-A*β*/DMPC, traj. 8) induces a significant compaction. The size and secondary structure of the peptide is, therefore, significantly modulated by the type of association when the latter occurs: electrostatic (strong interaction with bilayer surface) *versus* hydrophobic (penetration into the bilayer).

The penetration of the peptide into the membrane increases, as expected, the helix content. The maximal percentage of helix is displayed by the trajectories where the penetration is deeper: trajectory 1 for A*β*/DMPC and trajectory 8 for Cu-A*β*/DMPC, 15% and 20%, respectively (Table 4). These percentage is lower than that reported for A*β*(1-42) in micelles on the basis of CD and NMR experiments in SDS [78] and in helix-inducing solvents [79]. The difference is due to the partial achievement of peptide penetration in our simulation because of the exogenous addition. On the other hand, experiments in micelles and in apolar solvents are performed by peptide incoroporation into the microenvironment.

The number of intramolecular salt-bridges is, on average over the 10 trajectories, not altered in the presence of DMPC with respect to the case of water solution (Table 4). The SB quantity increases when the association of the peptide with DMPC is more significant (trajectories 1 and 5 for A*β*/DMPC, trajectory 8 for Cu-A*β*/DMPC). The number of contacts between positively charged groups in A*β* (see Methods) and P atoms, does not increase substantially, being always around 0.2, independently from the chosen trajectory (data not shown in Tables). The number of contacts between negatively charged groups in the peptide and the ammonium group in DMPC is always negligible, because of the steric effect of methyl groups attached to the N atom. These data indicate that the extent of association between peptide and membrane is independent from the electrostatic interactions between charged groups in the peptide and those with opposite charge at the membrane surface. Charged head groups in the membrane are on average not sufficient to divert charged groups in the peptide from pre-existent salt-bridges.

A further illustration of the type of interactions occurring in the peptide/DMPC association can be obtained by examining and comparing the final configurations of trajectories characterized by a different behavior. We limit this comparison, reported in Fig 10, to A*β*/DMPC, since the difference with Cu-A*β*/DMPC is, in this respect, marginal. The final configuration in trajectory 2 (top) represents a typical weak interaction between an almost unperturbed A*β* peptide and the surface of DMPC. Trajectory 5 (middle) ends with configurations significantly penetrating the membrane bilayer, but with interactions almost confined to the surface. Finally, in trajectory 1 the peptide rapidly achieves the penetration of the bilayer from the side of its C-terminus (bottom). In the latter conditions, it can be noticed that the region of A*β* crossing the layer surface is small, separating the N-terminus (above the surface) and the C-terminus (below the surface). This configuration, again, represents the requirement of removing the cross-talk between the N-terminus and the C-terminus (exerted by the bending of N-terminus towards the C-terminus) before a deeper penetration of the peptide into the membrane from the side of the C-terminus. This configuration is similar to that obtained by REMD of Cu-A*β*/DMPC displaying the deepest penetration into the bilayer (Fig 6B), with the main difference that the N-terminus is not partially neutralized by Cu binding.

**Fig 10.**
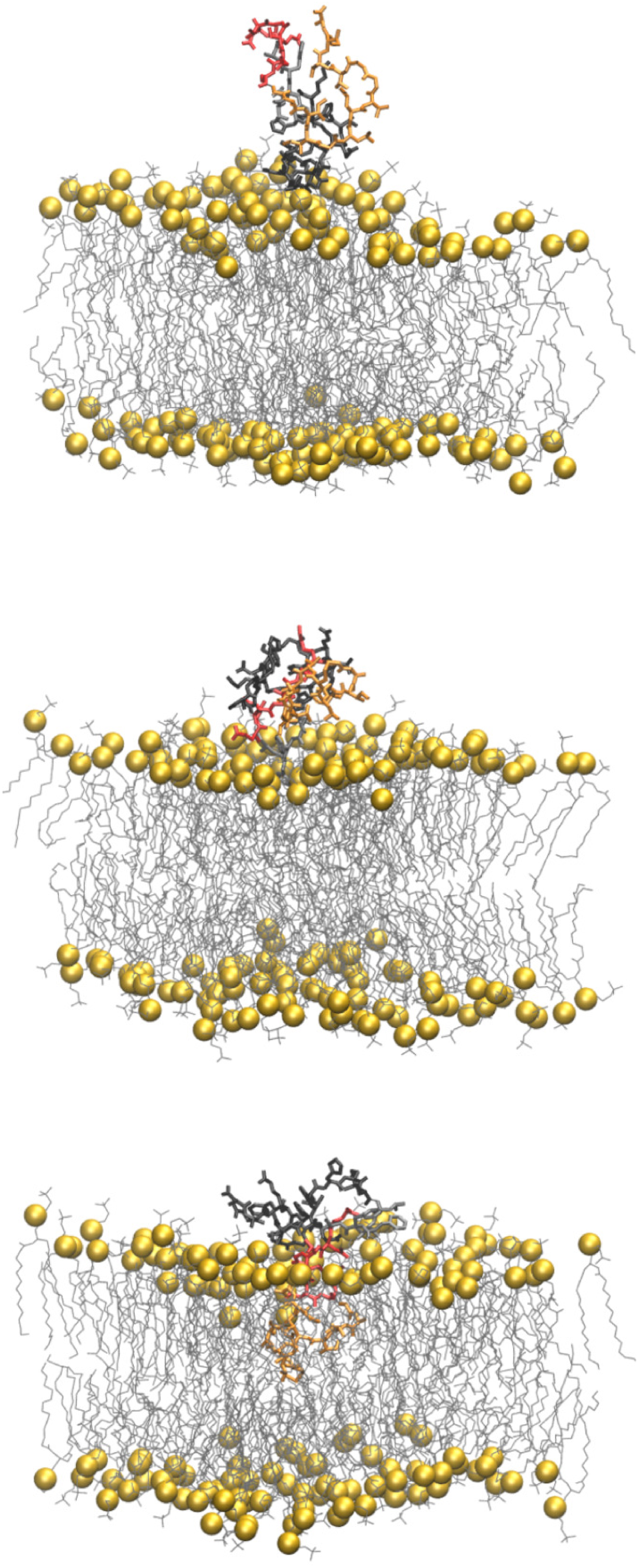
On the mechanism of penetration into DMPC. Final configurations of A*β*/DMPC in trajectories 2 (top), 5 (middle), and 1 (bottom). Residues 1-16 are in black (segment S1 in Fig 11), 17-21 in gray (S2), 22-28 in red (S3), and 29-42 in (S4). The peptide is represented as bond sticks. P atoms in DMPC are represented as yellow spheres. The other DMPC atoms are represented as lines. Water molecules and ions are not displayed. Bond and atomic radii are arbitrary.

The further comparison between statistical properties in the three different simulations represented by the snapshots described above, confirms the description of the force that is exerted by the DMPC bilayer when the peptide is incorporated. In the left panels of Fig 11 the probability of inter-residue contacts (see Methods) is displayed for trajectories 2 (top), 5 (middle), and 1 (bottom panels). In the first case there are almost no interactions between A*β*(1-42) and DMPC, since the number of A*β*-DMPC contacts is 5. In trajectory 5 significant interactions of A*β*(1-42) with the bilayer surface are revealed by an increase in the number of A*β*-DMPC contacts to 13. Finally, in trajectory 1 the deepest penetration of the peptide into the bilayer occurs and the number of contacts increases to 49. Again, trajectory 2 (top panel) displays a typical behavior for an unperturbed A*β*(1-42) peptide, where a weak cross-talk between many residues is allowed by the structural disorder of the peptide. As already observed for the monomeric A*β*(1-42) peptide in water solution, contacts are distributed among two domains, one N-terminal and one C-terminal, as it is shown by the low probability of contacts in the range of residues 20-26. In the case of interactions confined to the DMPC bilayer surface (trajectory 5, middle panel), we observe a conformational freezing, displayed by an increase, with respect to the free peptide, of highly populated contacts between residues far in the sequence. Some of them involve Glu 22, Asp 23, and Lys 28, with these charged sidechains interacting mostly with the N-terminus and not between themselves. In the case of a peptide that is more significantly embedded into the bilayer (trajectory 1, bottom panel), one notices the disappearing of contacts within residues in the C-terminus and the extension of the N-terminal domain up to Lys 28, with the void observed for trajectory 2 (top) almost filled. This change in cross-talk is induced by the formation of contacts between the C-terminus and DMPC. In the right panels of the same figure, we display the mass density for different atomic sets in A*β*(1-42). S1 is the N-terminus, S4 the C-terminus, while S2 is the hydrophobic segment and S3 contains the charged residues involved in one of the intramolecular SBs. When the peptide is out from the bilayer (trajectory 2, top-right panel) only the N-terminus (S1) is approaching the bilayer surface. The analysis of the trajectories not displaying penetration into the bilayer (all trajectories except 1 and 5, data not shown here) shows that there is no preference among the different segments for weak interactions with the bilayer surface. When a more significant interaction with the bilayer surface occurs (trajectory 5, middle-right panel) the hydrophobic segment S2 is projected towards the bilayer because of the stronger interactions among S3 and S1 (as shown in the middle-left panel). When the penetration is deeper (trajectory 1, bottom-right panel), the S4 segment overtakes the layer of P atoms, with the latter interacting with S3. Interestingly, in these conditions the S2 segment is projected towards the water layer, thus allowing interactions with other monomers in the nearby, especially if pre-organized as in trajectory 5 (middle panel).

**Fig 11.**
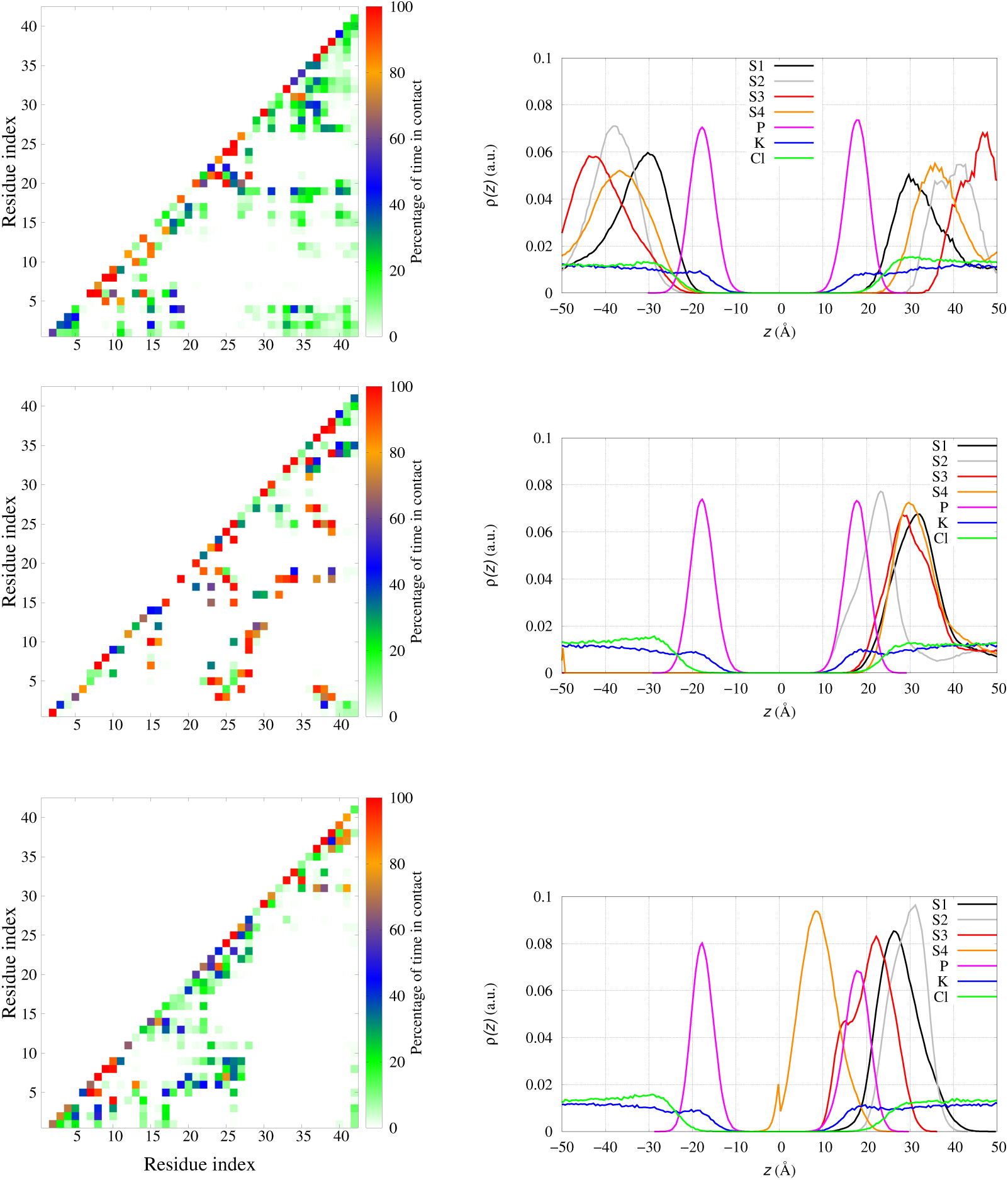
On the mechanism of penetration into DMPC. Probability, for A*β*/DMPC, of inter-residue contacts (left panels, see Methods for details) and density of mass for different atomic sets as a function of the coordinate *z* along the bilayer normal (right panels): trajectory 2 (top); trajectory 5 (middle); trajectory 1 (bottom). S1 are residues 1-16, S2 17-21, S3 22-28, S4 29-42. The density of each component is divided by the number of atoms in each atomic set.

As for Cu-A*β*/DMPC, 9 of 10 trajectories of display the behavior of A*β*/DMPC in trajectory 2, while only trajectory 8 displays a pattern similar to trajectory 5 in A*β*/DMPC.

The observations related to contacts, both defined as specific salt-bridges and generic inter-residue contacts, represent the process of changing the cross-talk between domains that are polymorphic in the free A*β*(1-42) peptide. The interactions with the charges on the surface of the membrane bilayer selects configurations that have a low population in the DMPC-unbound state, thus indicating a free energy barrier in the process of peptide penetration through the bilayer surface. The observation that peptide embedding into the membrane is a rare event (1 trajectory over 10) shows that the structural changes accompanying the penetration are hindered by the polymorphism that characterizes the monomeric A*β*(1-42) peptide. The Cu binding to A*β*(1-40) enhances the spread of configurations over polymorphic states in the monomeric state [61], thus providing a possible entropic explanation to the question why Cu binding reduces the penetration of monomers through the charged DMPC surface.

Again, we remind that this analysis is limited to peptide monomers.

## Discussion

In previous works we analyzed in detail the effect of Cu-binding on the properties of A*β*(1-42) peptide, both in monomeric and dimeric states. One major result of our models for monomers in water is that the interactions between the peptide charged sidechains and the water solvent are enhanced by Cu-binding. This observation is consistent with the longer life-time of Cu-A*β*(1-42) monomers compared to A*β*(1-42), when the Cu:A*β* ratio is 1:1, *i*.*e*. when all peptides are bound to Cu [80].

Therefore, by adding A*β*(1-42) and Cu-A*β*(1-42) monomers to DMPC, *i*.*e*. a lipid bilayer with charged head groups, the difference in organization of charged sidechains is potentially important.

Our models of monomeric A*β*(1-42) and Cu-A*β*(1-42) in contact with the DMPC bilayer confirm the experimental information that the exogenous addition to DMPC of these peptides reveals peptide-membrane interactions that are confined to the charged head groups of the bilayer. The interactions between the peptide and the membrane are concentrated in the head groups also in the few exceptions where the peptides are significantly embedded into the membrane bilayer. The exogenous addition of the peptide to the membrane bilayer does not alter significantly the bilayer structure when free divalent cations are either absent or bound to the peptide. As for the A*β*(1-42) peptide, this fact has been already observed experimentally by means of spectroscopy and diffraction studies [81]. Consistently, dramatic changes of peptide/membrane interactions are observed at conditions where the peptide is truncated to be more hydrophobic (A*β*(25-35)) or forms fibril assemblies [81, 82].

The picture of A*β* monomers floating over the membrane surface is consistent with other observations reported in the literature. A recent FRET experimental work [4] describes the strong interactions among growing fibrils and the DOPC membrane, modelled as a lipid vesicle. The same study confirms that monomers do not directly bind the lipid bilayer, as already observed in previous studies.

As for the impact on oligomer formation, our results point out the possible role of charged groups of the bilayer in organizing monomers into oligomers. Indeed, several simulations showed that a strong association between A*β* and zwitterionic and charged membranes occurs starting from tetrameric A*β* assemblies [15]. Since it is known that the lag-time of monomers associated to Cu is larger than that of Cu-free A*β* when in the water solvent [80], it is not surprising that the DMPC association with A*β*(1-42) in the absence of divalent cations does not decrease the chance of inter-monomer contacts compared to the water solution. The bilayer-water interface, when bilayer has charged groups on the surface, exerts a mild attraction for A*β*(1-42) thus decreasing the freedom of monomers by reducing the space dimensionality. Conversely, at oligomeric level the bilayer surface can assist the formation of larger oligomers and protofibrils. This type of association has been observed in models of preformed protofibrils interacting with lipid bilayers [14].

We notice here that in ss-NMR experiments, the effect of the addition of free Cu^2+^ and Zn^2+^ ions on the membrane properties is more dramatic than in the presence of A*β* peptide [53]. Similar strong effects have been observed both experimentally and computationally for free Ca^2+^ ions [41–43], and Mg and Cu divalent cations are even smaller than Ca^2+^ in size. For the first time, we show in this study that the A*β*-bound Cu^2+^ ion does not exert the strong perturbation on the membrane exerted by a free divalent ion. Indeed, the effect of Cu-A*β* monomer on the membrane is weaker than that of the more charged A*β* peptide.

Therefore, the formation of the Cu-A*β* complex before an eventual incorporation into the membrane and before an increase in peptide concentration, appears as a protection against membrane destabilization and oxidation. This hypothesis is confirmed by the NMR experiments performed with the A*β*(25-35) peptide, both without and with Cu [54, 81]. Since the N-truncated peptide does not bind Cu, the addition of Cu to the system has an effect on the bilayer that is similar to that of free Cu.

## Conclusion

We perturbed an atomistic model of the DMPC bilayer, representing a very crude approximation of a portion of the cellular membrane facing the synaptic region between neurons, with a single divalent cation (Mg^2+^) and with Cu-free and Cu-loaded amyloid-*β* peptides of 42 aminoacid residues in the monomeric form.

All the data reported in our simulations represent important structural and electrostatic changes of the bilayer when a single divalent cation interacts with the phosphate groups of DMPC. On the other hand, the presence of the peptide represents a floating molecule, mildly interacting with the bilayer surface, and well suited to sequester divalent cations, in this case Cu^2+^. The model clearly depicts the possible protective role of the amyloid-*β*(1-42) peptide in avoiding interactions between Cu^2+^ and the synaptic membrane.

The model has many limitations. Beyond the limitations in size and number of components, that are common to application of atomistic models, there is the lack of working approximations to interactions between a ion like Cu^2+^, with available 3d orbitals, and molecules providing a pletora of possible ligand atoms, like phosphate, carboxylate, imidazole, and carbonyl groups, not to mention deprotonated amide backbone nitrogen that are known to bind Cu^2+^ at physiological pH. These limitations will be removed by polarizable and reactive force-fields, not yet available.

The investigation of the events occurring when the concentration of the peptide increases are the future perspective of this study. However, the type of weak interactions of the peptide with DMPC, shows that modulation of inter-peptide electrostatic interactions are likely changing the picture describing the behavior of monomers, where intramolecular salt-bridges are found particularly stable. The assembly of several monomers into oligomers, especially when loaded with Cu^2+^, is likely affecting the surface of the bilayer. Then, as expected, the increase in concentration of Cu-A*β*(1-42) in the synaptic region becomes the crucial event destabilizing the neuron membrane. The increase in turnover of Cu-A*β* monomers or dimers, possibly because of self-oxidation (the latter enhanced in dimers), can also contribute to membrane protection.

## Supporting information

**S1 File. US**. Description of umbrella sampling estimate of binding free energy to DMPC for Mg^2+^ and Na^+^.

**S2 File. DMPC**. Initial coordinates for DMPC MD simulations (1-4). No water molecules and KCl are included. Format is PDB for first configuration and compressed (Bzip2) XYZ format for the other configurations.

**S3 File. Mg/DMPC**. Initial coordinates for Mg/DMPC MD simulations (1-3). No water molecules and KCl are included. Format is the same as for DMPC.

**S4 File. A***β***/DMPC**. Initial coordinates for A*β*/DMPC CMD simulations (1-10). No water molecules and KCl are included. Format is the same as for DMPC.

**S5 File. Cu-A***β***/DMPC**. Initial coordinates for Cu-A*β*/DMPC CMD simulations (1-10). No water molecules and KCl are included. Format is the same as for DMPC.

**S6 File. A***β***/DMPC**. Initial coordinates for A*β*/DMPC REMD simulations (1-56). No water molecules and KCl are included. Format is the same as for DMPC.

**S7 File. Cu-A***β***/DMPC**. Initial coordinates for Cu-A*β*/DMPC REMD simulations (1-56). No water molecules and KCl are included. Format is the same as for DMPC.

## Acknowledgments

The work was supported by: Narodowe Centrum Nauki in Poland (grant n. 2015/19/B/ST4/02721); Department of Science and Technology (Ho Chi Minh City, Vietnam); PLGrid Infrastructure (Poland); the bilateral project Cnr(I)-PAN(PL) “The role of copper ions in neurodegeneration: molecular models”.

## References

1. Blennow K, de Leon MJ, Zetterberg H. Alzheimer’s disease. Lancet. 2006;368(9533):387–403. doi: 10.1016/S0140-6736(06)69113-7.

2. Müller UC, Deller T. Editorial: The Physiological Functions of the APP Gene Family. Front Mol Neurosci. 2017;10:334. doi: 10.3389/fnmol.2017.00334.

3. Kummer MP, Heneka MT. Truncated and modified amyloid-beta species. Alzheimers Res Ther. 2014;6(3):28. doi: 10.1186/alzrt258.

4. Lindberg DJ, Wesén E, Björkeroth J, Rocha S, Esbjörner EK. Lipid membranes catalyse the fibril formation of the amyloid-*β* (1-42) peptide through lipid-fibril interactions that reinforce secondary pathways. Biochim Biophys Acta, Biomembr. 2017;1859(10):1921–1929. doi: 10.1016/j.bbamem.2017.05.012.

5. Mucke L, Masliah E, Yu GQ, Mallory M, Rockenstein EM, Tatsuno G, et al. High-Level Neuronal Expression of A*β*1–42 in Wild-Type Human Amyloid Protein Precursor Transgenic Mice: Synaptotoxicity without Plaque Formation. J Neurosci. 2000;20(11):4050–4058. doi: 10.1523/JNEUROSCI.20-11-04050.2000.

6. Evangelisti E, Cascella R, Becatti M, Marrazza G, Dobson CM, Chiti F, et al. Binding affinity of amyloid oligomers to cellular membranes is a generic indicator of cellular dysfunction in protein misfolding diseases. Sci Rep. 2016;6:32721. doi: 10.1038/srep32721.

7. Niu Z, Zhang Z, Zhao W, Yang J. Interactions between amyloid-*β* peptide and lipid membranes. Biochim Biophys Acta, Biomembr. 2018;1860(9):1663–1669. doi: 10.1016/j.bbamem.2018.04.004.

8. Kepp KP. Alzheimer’s disease due to loss of function: A new synthesis of the available data. Prog Neurobiol. 2016;143:36–60. doi: 10.1016/j.pneurobio.2016.06.004.

9. Marrink SJ, Corradi V, Souza PCT, Ingólfsson HI, Tieleman DP, Sansom MSP. Computational Modeling of Realistic Cell Membranes. Chem Rev. 2019;119(9):6184–6226. doi: 10.1021/acs.chemrev.8b00460.

10. Lemkul JA, Bevan DR. A comparative molecular dynamics analysis of the amyloid *β*-peptide in a lipid bilayer. Arch Biochem Biophys. 2008;470(1):54–63. doi: 10.1016/j.abb.2007.11.004.

11. Lockhart C, Klimov DK. Alzheimer’s A*β*10-40 Peptide Binds and Penetrates DMPC Bilayer: An Isobaric-Isothermal Replica Exchange Molecular Dynamics Study. J Phys Chem B. 2014;118(10):2638–2648. doi: 10.1021/jp412153s.

12. Friedman R, Pellarin R, Caflisch A. Amyloid Aggregation on Lipid Bilayers and Its Impact on Membrane Permeability. J Mol Biol. 2009;387(2):407–415. doi: 10.1016/j.jmb.2008.12.036.

13. Liu L, Hyeon C. Contact Statistics Highlight Distinct Organizing Principles of Proteins and RNA. Biophys J. 2016;110(11):2320–2327. doi: 10.1016/j.bpj.2016.04.020.

14. Tofoleanu F, Buchete NV. Molecular Interactions of Alzheimer’s A*β* Protofilaments with Lipid Membranes. J Mol Biol. 2012;421(4-5):572–586. doi: 10.1016/j.jmb.2011.12.063.

15. Poojari C, Kukol A, Strödel B. How the amyloid-beta peptide and membranes affect each other: An extensive simulation study. Biochim Biophys Acta, Biomembr. 2013;1828(2):327–339. doi: 10.1016/j.bbamem.2012.09.001.

16. Tofoleanu F, Brooks BR, Buchete NV. Modulation of Alzheimer’s A*β* Protofilament-Membrane Interactions by Lipid Headgroups. ACS Chem Neurosci. 2015;6:446–455. doi: 10.1021/cn500277f.

17. Brown A, Bevan D. Molecular Dynamics Simulations of Amyloid *β*-Peptide(1-42): Tetramer Formation and Membrane Interactions. Biophys J. 2016;111(5):937–949. doi: 10.1016/j.bpj.2016.08.001.

18. Press-Sandler O, Miller Y. Molecular mechanisms of membrane-associated amyloid aggregation: Computational perspective and challenges. Biochim Biophys Acta, Biomembr. 2018;1860(9):1889–1905. doi: 10.1016/j.bbamem.2018.03.014.

19. Friedman R. Membrane–Ion Interactions. J Membr Biol. 2018;251(3):453–460. doi: 10.1007/s00232-017-0010-y.

20. Ackerman CM, Chang CJ. Copper signaling in the brain and beyond. J Biol Chem. 2018;293(13):4628–4635. doi: 10.1074/jbc.R117.000176.

21. Hartter DE, Barnea A. Evidence for release of copper in the brain: Depolarization-induced release of newly taken-up 67copper. Synapse. 1988;2(4):412–415. doi: 10.1002/syn.890020408.

22. Vassallo N, Herms J. Cellular prion protein function in copper homeostasis and redox signalling at the synapse. J Neurochem. 2003;86(3):538–544. doi: 10.1046/j.1471-4159.2003.01882.x.

23. Wild K, August A, Pietrzik CU, Kins S. Structure and Synaptic Function of Metal Binding to the Amyloid Precursor Protein and its Proteolytic Fragments. Front Mol Neurosci. 2017;10:21. doi: 10.3389/fnmol.2017.00021.

24. Rae TD, Schmidt PJ, Pufahl RA, Culotta VC, O’Halloran TV. Undetectable Intracellular Free Copper: The Requirement of a Copper Chaperone for Superoxide Dismutase. Science. 1999;284(5415):805–808. doi: 10.1126/science.284.5415.805.

25. Kepp KP. Alzheimer’s disease: How metal ions define *β*-amyloid function. Coord Chem Rev. 2017;351:127–159. doi: 10.1016/j.ccr.2017.05.007.

26. Multhaup G, Schlicksupp A, Hesse L, Beher D, Ruppert T, Masters CL, et al. The Amyloid Precursor Protein of Alzheimer’s Disease in the Reduction of Copper(II) to Copper(I). Science. 1996;271(5254):1406–1409. doi: 10.1126/science.271.5254.1406.

27. Strausak D, Mercer JFB, Dieter HH, Stremmel W, Multhaup G. Copper in disorders with neurological symptoms: Alzheimer’s, Menkes, and Wilson diseases. Brain Res Bull. 2001;55(2):175–185. doi: 10.1016/S0361-9230(01)00454-3.

28. Gaggelli E, Kozlowski H, Valensin D, Valensin G. Copper Homeostasis and Neurodegenerative Disorders (Alzheimer’s, Prion, and Parkinson’s Diseases and Amyotrophic Lateral Sclerosis). Chem Rev. 2006;106(6):1995–2044. doi: 10.1021/cr040410w.

29. Ohba S, Hiramatsu M, Edamatsu R, Mori I, Mori A. Metal ions affect neuronal membrane fluidity of rat cerebral cortex. Neurochem Res. 1994;19(3):237–241. doi: 10.1007/BF00971570.

30. Suwalsky M, Ungerer B, Quevedo L, Aguilar F, Sotomayor CP. Cu2+ ions interact with cell membranes. J Inorg Biochem. 1998;70(3):233–238. doi: 10.1016/S0162-0134(98)10021-1.

31. García JJ, Martínez-Ballarín E, Millán-Plano S, Allué JL, Albendea C, Fuentes L, et al. Effects of trace elements on membrane fluidity. J Trace Elem Med Biol. 2005;19(1):19–22. doi: 10.1016/j.jtemb.2005.07.007.

32. Jiang X, Zhang J, Zhou B, Li P, Hu X, Zhu Z, et al. Anomalous behavior of membrane fluidity caused by copper-copper bond coupled phospholipids. Sci Rep. 2018;8(1):14093–15002. doi: 10.1038/s41598-018-32322-4.

33. Quist A, Doudevski I, Lin H, Azimova R, Ng D, Frangione B, et al. Amyloid ion channels: A common structural link for protein-misfolding disease. Proc Natl Acad Sci USA. 2005;102(30):10427–10432. doi: 10.1073/pnas.0502066102.

34. Di Scala C, Chahinian H, Yahi N, Garmy N, Fantini J. Interaction of Alzheimer’s *β*-Amyloid Peptides with Cholesterol: Mechanistic Insights into Amyloid Pore Formation. Biochemistry. 2014;53(28):4489–4502. doi: 10.1021/bi500373k.

35. Reybier K, Ayala S, Alies B, Rodrigues JaV, Bustos Rodriguez S, La Penna G, et al. Free superoxide is an intermediate in the production of H2O2 by copper(I)-A*β* peptide and O2. Angew Chem Intl Ed. 2016;55:1085–1089. doi: 10.1002/anie.201508597.

36. La Penna G, Mai Suan L. Towards a high-throughput modelling of copper reactivity induced by structural disorder in amyloid peptides. Chem Eur J. 2018;24:5259–5270. doi: 10.1002/chem.201704654.

37. Lynch T, Cherny RA, Bush AI. Oxidative processes in Alzheimer’s disease: the role of A*β*-metal interactions. Experim Gerontol. 2000;35(4):445–451. doi: 10.1016/S0531-5565(00)00112-1.

38. Perry G, Sayre LM, Atwood CS, Castellani RJ, Cash AD, Rottkamp CA, et al. The Role of Iron and Copper in the Aetiology of Neurodegenerative Disorders. CNS Drugs. 2002;16(5):339–352. doi: 10.2165/00023210-200216050-00006.

39. Bagheri S, Squitti R, Haertlé T, Siotto M, Saboury AA. Role of Copper in the Onset of Alzheimer’s Disease Compared to Other Metals. Front Aging Neurosci. 2018;9:446. doi: 10.3389/fnagi.2017.00446.

40. Widomska J, Raguz M, Subczynski WK. Oxygen permeability of the lipid bilayer membrane made of calf lens lipids. Biochim Biophys Acta, Biomembr. 2007;1768(10):2635–2645. doi: 10.1016/j.bbamem.2007.06.018.

41. Javanainen M, Melcrová A, Magarkar A, Jurkiewicz P, Hof M, Jungwirth P, et al. Two cations, two mechanisms: interactions of sodium and calcium with zwitterionic lipid membranes. Chem Commun. 2017;53(39):5380–5383. doi: 10.1039/C7CC02208E.

42. Bilkova E, Pleskot R, Rissanen S, Sun S, Czogalla A, Cwiklik L, et al. Calcium Directly Regulates Phosphatidylinositol 4,5-Bisphosphate Headgroup Conformation and Recognition. J Am Chem Soc. 2017;139(11):4019–4024. doi: 10.1021/jacs.6b11760.

43. Melcr J, Martinez-Seara H, Nencini R, Kolafa J, Jungwirth P, Ollila OHS. Accurate Binding of Sodium and Calcium to a POPC Bilayer by Effective Inclusion of Electronic Polarization. J Phys Chem B. 2018;122(16):4546–4557. doi: 10.1021/acs.jpcb.7b12510.

44. Nguyen HT, Hori N, Thirumalai D. Theory and simulations for RNA folding in mixtures of monovalent and divalent cations. Proc Natl Acad Sci. 2019;116(42):21022–21030. doi: 10.1073/pnas.1911632116.

45. Miller Y, Ma B, Nussinov R. Metal binding sites in amyloid oligomers: Complexes and mechanisms. Coord Chem Rev. 2012;256(19-20):2245–2252. doi: 10.1016/j.ccr.2011.12.022.

46. Wineman-Fisher V, Bloch DN, Miller Y. Challenges in studying the structures of metal-amyloid oligomers related to type 2 diabetes, Parkinson’s disease, and Alzheimer’s disease. Coord Chem Rev. 2016;327-328:20–26. doi: 10.1016/j.ccr.2016.04.010.

47. Duarte F, Bauer P, Barrozo A, Amrein BA, Purg M, Åqvist J, et al. Force Field Independent Metal Parameters Using a Nonbonded Dummy Model. J Phys Chem B. 2014;118(16):4351–4362. doi: 10.1021/jp501737x.

48. La Penna G, Chelli R. Structural Insights into the Osteopontin-Aptamer Complex by Molecular Dynamics Simulations. Front Chem. 2018;6:1–11. doi: 10.3389/fchem.2018.00002.

49. Drew SC, Masters CL, Barnham KJ. Alanine-2 carbonyl is an oxygen ligand in Cu2+ coordination of Alzheimer’s disease amyloid-*β* peptide: Relevance to N-terminally truncated forms. J Am Chem Soc. 2009;131(21):8760–8761. doi: 10.1021/ja903669a.

50. Dorlet P, Gambarelli S, Faller P, Hureau C. Pulse EPR spectroscopy reveals the coordination sphere of copper(II) ions in the 1-16 amyloid-*β* peptide: A key role of the first two N-terminus residues. Angew Chem Int Ed. 2009;48(49):9273–9276. doi: 10.1002/anie.200904567.

51. Kim D, Kim NH, Kim SH. 34 GHz Pulsed ENDOR Characterization of the Copper Coordination of an Amyloid *β* Peptide Relevant to Alzheimer’s Disease. Angew Chem Intl Ed. 2013;52(4):1139–1142. doi: 10.1002/anie.201208108.

52. Huy PDQ, Vuong QV, La Penna G, Faller P, Li MS. Impact of Cu(II) binding on structures and dynamics of A*β*42 monomer and dimer: Molecular dynamics study. ACS Chem Neurosci. 2016;7(10):1348–1363. doi: 10.1021/acschemneuro.6b00109.

53. Lau TL, Ambroggio EE, Tew DJ, Cappai R, Masters CL, Fidelio GD, et al. Amyloid-*β* Peptide Disruption of Lipid Membranes and the Effect of Metal Ions. J Mol Biol. 2006;356(3):759–770. doi: 10.1016/j.jmb.2005.11.091.

54. Lau TL, Gehman JD, Wade JD, Perez K, Masters CL, Barnham KJ, et al. Membrane interactions and the effect of metal ions of the amyloidogenic fragment A*β*(25-35) in comparison to A*β*(1-42). Biochim Biophys Acta, Biomembr. 2007;1768(10):2400–2408. doi: 10.1016/j.bbamem.2007.05.004.

55. Goldberg M, De Pittà M, Volman V, Berry H, Ben-Jacob E. Nonlinear Gap Junctions Enable Long-Distance Propagation of Pulsating Calcium Waves in Astrocyte Networks. PLOS Comput Biol. 2010;6(8):1–14. doi: 10.1371/journal.pcbi.1000909.

56. Case DA, Betz RM, Cerutti DS, Cheatham III TE, Darden TA, E DR, et al. AMBER 2016. San Francisco, USA: University of California at San Fransisco; 2016.

57. Maier JA, Martinez C, Kasavajhala K, Wickstrom L, Hauser KE, Simmerling C. ff14SB: Improving the Accuracy of Protein Side Chain and Backbone Parameters from ff99SB. J Chem Theory Comput. 2015;11(8):3696–3713. doi: 10.1021/acs.jctc.5b00255.

58. Jorgensen WL, Chandrasekhar J, Madura JD, Impey RW, Klein MJ. Comparison of simple potential functions for simulating liquid water. J Chem Phys. 1983;79:926–935. doi: 10.1063/1.445869.

59. Dickson CJ, Madej BD, Skjevik ÅA, Betz RM, Teigen K, Gould IR, et al. Lipid14: The Amber Lipid Force Field. J Chem Theory Comput. 2014;10(2):865–879. doi: 10.1021/ct4010307.

60. Hornak V, Abel R, Okur A, Strockbine B, Roitberg A, Simmerling C. Comparison of multiple Amber force fields and development of improved protein backbone parameters. Proteins: Struct, Funct, Bioinf. 2006;65(3):712–725. doi: 10.1002/prot.21123.

61. Pham DQH, Li MS, La Penna G. Copper binding induces polymorphism in amyloid-*β* peptide: Results of computational models. J Phys Chem B. 2018;122(29):7243–7252. doi: 10.1021/acs.jpcb.8b03983.

62. Carballo-Pacheco M, Strodel B. Comparison of force fields for Alzheimer’s A*β*42: A case study for intrinsically disordered proteins. Protein Sci. 2017;26(2):174–185. doi: 10.1002/pro.3064.

63. Man VH, He X, Derreumaux P, Ji B, Xie XQ, Nguyen PH, et al. Effects of All-Atom Molecular Mechanics Force Fields on Amyloid Peptide Assembly: The Case of A*β*(16-22) Dimer. J Chem Theory Comput. 2019;15(2):1440–1452. doi: 10.1021/acs.jctc.8b01107.

64. Krupa P, Quoc Huy PD, Li MS. Properties of monomeric A*β*42 probed by different sampling methods and force fields: Role of energy components. J Chem Phys. 2019;151(5):055101. doi: 10.1063/1.5093184.

65. Bayly CI, Cieplak P, Cornell W, Kollman PA. A Well-behaved Electrostatic Potential Based Method Using Charge Restraints for Deriving Atomic Charges: the RESP Model. J Phys Chem. 1993;97(40):10269–10280. doi: 10.1021/j100142a004.

66. Cornell WD, Cieplak P, Bayly CI, Kollman PA. Application of RESP charges to calculate conformational energies, hydrogen bond energies, and free energies of solvation. J Am Chem Soc. 1993;115(21):9620–9631. doi: 10.1021/ja00074a030.

67. Comba P, Remenyi R. A new molecular mechanics force field for the oxidized form of blue copper proteins. J Comput Chem. 2002;23(7):697–705. doi: 10.1002/jcc.10084.

68. Wu XW, Brooks BR. Self-guided langevin dynamics simulation method. Chem Phys Lett. 2003;381(3-4):512–518. doi: 10.1016/j.cplett.2003.10.013.

69. Ryckaert JP, Ciccotti G, Berendsen HJC. Numerical integration of the cartesian equations of motion with constraints: Molecular dynamics of n-alkanes. J Comput Phys. 1977;23:327–341. doi: 10.1016/0021-9991(77)90098-5.

70. Darden T, York D, Pedersen L. Particle mesh Ewald: An N-log(N) method for Ewald sums in large systems. J Chem Phys. 1993;98:10089–10092. doi: 10.1063/1.464397.

71. Weiser J, Shenkin PS, Still WC. Approximate atomic surfaces from linear combinations of pairwise overlaps (LCPO). J Comput Chem. 1999;20(2):217–230. doi: 10.1002/(SICI)1096-987X(19990130)20:2<217::AID-JCC4>3.0.CO;2-A.

72. Levine ZA, Venable RM, Watson MC, Lerner MG, Shea JE, Pastor RW, et al. Determination of Biomembrane Bending Moduli in Fully Atomistic Simulations. J Am Chem Soc. 2014;136(39):13582–13585. doi: 10.1021/ja507910r.

73. Watson MC, Penev ES, Welch PM, Brown FLH. Thermal fluctuations in shape, thickness, and molecular orientation in lipid bilayers. J Chem Phys. 2011;135(24):244701. doi: 10.1063/1.3660673.

74. Marcus Y. Thermodynamics of solvation of ions. Part 5. Gibbs free energy of hydration at 298.15 K. J Chem Soc, Faraday Trans. 1991;87:2995–2999. doi: 10.1039/FT9918702995.

75. Kučerka N, Nieh MP, Katsaras J. Fluid phase lipid areas and bilayer thicknesses of commonly used phosphatidylcholines as a function of temperature. Biochim Biophys Acta, Biomembr. 2011;1808(11):2761–2771. doi: https://doi.org/10.1016/j.bbamem.2011.07.022.

76. López CA, Unkefer CJ, Swanson BI, Swanson JMJ, Gnanakaran S. Membrane perturbing properties of toxin mycolactone from Mycobacterium ulcerans. PLOS Comput Biol. 2018;14(2):e1005972–22. doi: 10.1371/journal.pcbi.1005972.

77. Jarvet J, Danielsson J, Damberg P, Oleszczuk M, Gräslund A. Positioning of the Alzheimer A*β*(1-40) peptide in SDS micelles using NMR and paramagnetic probes. J Biomol NMR. 2007;39(1):63–72. doi: 10.1007/s10858-007-9176-4.

78. Shao H, Jao Sc, Ma K, Zagorski MG. Solution structures of micelle-bound amyloid *β*-(1-40) and *β*-(1-42) peptides of Alzheimer’s disease. J Mol Biol. 1999;285(2):755–773. doi: 10.1006/jmbi.1998.2348.

79. Crescenzi O, Tomaselli S, Guerrini R, Salvadori S, D’Ursi AM, Temussi PA, et al. Solution structure of the Alzheimer amyloid *β*-peptide (1-42) in an apolar microenvironment. Eur J Biochem. 2002;269(22):5642–5648. doi: 10.1046/j.1432-1033.2002.03271.x.

80. Pedersen JT, Østergaard J, Rozlosnik N, Gammelgaard B, Heegaard NHH. Cu(II) mediates kinetically distinct, non-amyloidogenic aggregation of amyloid-*β* peptide. J Biol Chem. 2011;286(30):26952–26963. doi: 10.1074/jbc.M111.220863.

81. Accardo A, Shalabaeva V, Cotte M, Burghammer M, Krahne R, Riekel C, et al. Amyloid *β* Peptide Conformational Changes in the Presence of a Lipid Membrane System. Langmuir. 2014;30(11):3191–3198. doi: 10.1021/la500145r.

82. Kandel N, Zheng T, Huo Q, Tatulian SA. Membrane Binding and Pore Formation by a Cytotoxic Fragment of Amyloid *β* Peptide. J Phys Chem B. 2017;121(45):10293–10305. doi: 10.1021/acs.jpcb.7b07002.

